# A compact viral IRES translates a downstream open reading frame

**DOI:** 10.1101/2025.05.13.653779

**Authors:** Madeline E. Sherlock, Katherine E. Segar, Jeffrey S. Kieft

**Affiliations:** New York Structural Biology Center, New York, NY 10025, USA; Department of Biochemistry and Molecular Genetics, University of Colorado Anschutz Medical Campus, Aurora, Colorado, 80045, USA; Department of Biochemistry and Molecular Biophysics, Columbia University, New York, NY, USA; Department of Molecular, Cell and Developmental Biology, University of California, Santa Cruz, CA, USA

**Keywords:** internal ribosome entry site, viral RNA, cryoEM, translation regulation, RNA structure

## Abstract

Hepatitis C virus (HCV) and many other RNA viruses contain a Type IV internal ribosome entry site (IRES) in their 5′ untranslated region (UTR). These IRES RNAs adopt a complex tertiary structure that interacts directly with the ribosome, enabling cap-independent translation initiation. Using bioinformatic methods to search viral genomes for more Type IV IRES RNAs, we discovered that megrivirus E (MeV-E) contains a putative Type IV IRES within its annotated 3′ UTR. In addition to its unusual location downstream of the main coding region, the size of the MeV-E 3′ IRES is substantially reduced compared to known Type IV IRESs. We confirmed the secondary structure of the MeV-E 3′ IRES, determined its 3D structure in complex with the ribosome using cryoEM, and showed that the MeV-E 3′ IRES initiates translation but at lower levels compared to the larger Type IV IRES in the MeV-E 5′ UTR. We hypothesize that the absence of several domains in the MeV-E 3′ IRES compared to other Type IV IRESs results in a loss of ribosomal protein interactions and a relative decrease in translation activity. This small Type IV IRES enables translation of a second open reading frame in the MeV-E genome, which likely encodes a transmembrane protein that is conserved in other megriviruses. We propose a model wherein MeV-E expresses lower levels of its downstream-encoded protein compared to those in the upstream coding region using a pared down IRES structure, demonstrating purposeful tuning of translation through RNA structural variation.

## INTRODUCTION

Protein expression in eukaryotes is tightly regulated, especially at the initiation step, to ensure production of only the intended gene product. Canonical eukaryotic translation initiation depends on the 7-methylguanosine (m7G) cap at the 5’ end of the messenger RNA (mRNA) and requires many initiation factor proteins in a multistep process to assemble a ribosome at the authentic start codon.^1–3^ Most mature eukaryotic mRNAs are mono-cistronic, encoding only one protein product in the main open reading frame (ORF). In contrast, many eukaryotic viruses have multi-cistronic mRNAs and induce alternative modes of translation to produce the mature protein products necessary to proliferate.^4–8^ Viruses use numerous strategies to bypass canonical translation initiation requirements and decode regions that would not otherwise yield protein. Many such noncanonical translation events involve structured RNA elements embedded within the viral genome.^9^ For positive-sense, single-stranded RNA (+ssRNA) viruses the genome itself serves as the viral mRNA.

Internal ribosome entry site (IRES) RNAs are a widespread tactic employed by viruses to recruit ribosomes to start codons and initiate translation in a 5′ end- and m7G cap-independent manner.^10,11^ Multiple classes of IRESs, currently categorized into six different IRES types,^12^ have been identified in a range of viruses with disparate host specificity. Each type of IRES has a distinct RNA structural architecture along with varying eukaryotic initiation factor (eIF) requirements and ribosomal interactions.^10,12^

Hepatitis C virus (HCV) contains a Type IV IRES within the 5′ untranslated region (UTR) of its +ssRNA genome.^13–16^ The HCV IRES RNA comprises multiple stem loop domains, a four-way junction, two pseudoknots, and other tertiary structure features to enable direct binding to the small (40S) ribosomal subunit and the eIF3 complex.^17–27^ The HCV IRES structure and initiation mechanism have been extensively characterized^28–35^ and it serves as the prototypical Type IV IRES. Of the four stem loop structures in the HCV 5′ UTR, domains II and III (Fig. S1) are necessary for the IRES to initiate translation. Domain II reaches into the E-site of the ribosome and helps place the AUG start codon in the P-site.^22,30,36^ The “core region” of the HCV IRES centers around a four-way junction comprising helices at the base of domain III as well as stem loops IIId, IIIe, and IIIf (Fig. S1).^19,21,37,38^ Domain IIId serves as a key site of interaction with the ribosome, forming intermolecular base pairs with solvent-accessible nucleotides of the 18S ribosomal RNA (rRNA).^21,31,39^ Domains IIIa and IIIc interact with a ribosomal protein in the 40S subunit and, along with the apical domain IIIb, proteins in the eIF3 complex.^21,23,24,40^

While HCV serves as the archetype for Type IV IRESs, there is significant diversity in both sequence and structure within this class.^13,37^ Notably, some Type IV IRESs have large insertions or deletions, the functional implications of which are largely uncharacterized. To date, most Type IV IRESs have been identified by sequence gazing within viral 5′ UTRs or querying databases using sequence similarity.^37,41^

We suspected that more RNA structures belonging to the Type IV IRES class had yet to be identified because some aspect of their sequence, structure, viral phylogeny, or genomic context caused them to be overlooked. We employed a bioinformatic approach using an alignment of Type IV IRES structures, not primary sequence conservation alone, as an input model to search viral genomes for more examples. Herein, we characterize the structure and function of a new Type IV IRES, unique in its minimized domain architecture and genomic location, then discuss the broader implications of its discovery for regulating gene expression through RNA structure.

## RESULTS AND DISCUSSION

### Bioinformatic identification of a Type IV IRES in an unusual genomic location

We created a preliminary “seed” alignment of six Type IV (also previously referred to as class III) IRES RNAs, guided by previously determined secondary structure models. This alignment served as a structural model to inform the program Infernal^42,43^ to search a database of sequenced viral genomes for homologous IRES structures. Figure 1A contains an overview of the search strategy, which has been successful in expanding the known members of numerous other families of RNA structures.^44–50^ This homology search returned any sequence matching our input secondary structural model for Type IV IRES above a set significance threshold, with output files containing the aligned RNA sequences, their virus of origin, and their location within the viral genome.^42^ This enabled us to evaluate the phylogeny as well as genomic context of each putative Type IV IRES RNA. We report a detailed classification of all Type IV IRESs identified by this bioinformatic search in a complementary publication.^51^

**FIGURE 1.**
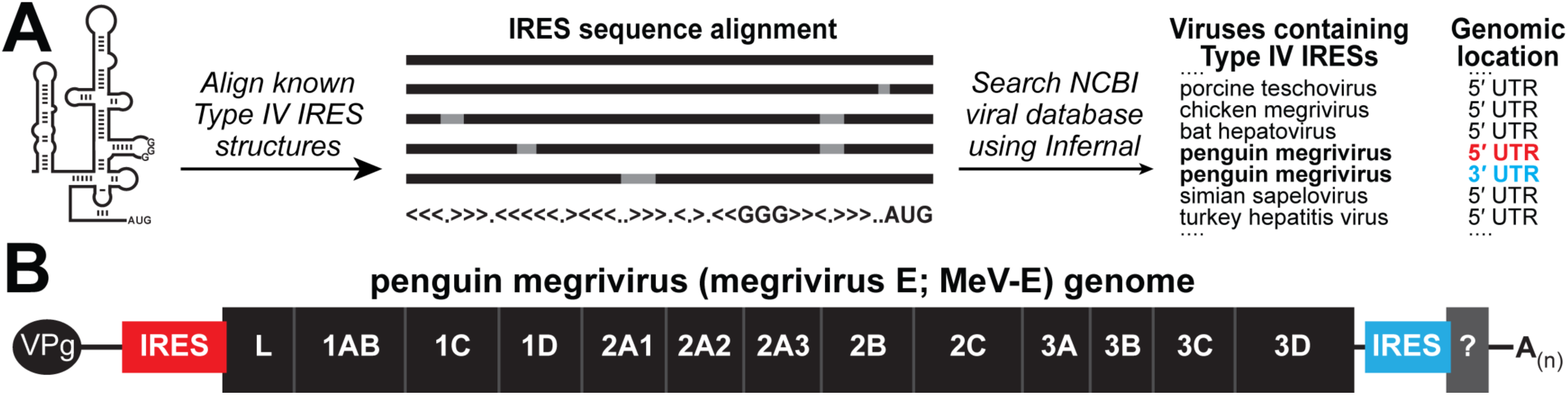
Bioinformatic identification of a Type IV IRES in a novel genomic location. (A) Overview of the bioinformatic homology-based search using an RNA structural alignment to identify Type IV IRESs, with the virus and genomic location of selected identified sequences displayed to the right. (B) Cartoon representation of the genome of penguin-infecting megrivirus E (MeV-E). This virus contains a Type IV IRES in its 5’ UTR (red) to express the main open reading frame that encodes the viral polyprotein, and a putative IRES in the region previously annotated as the 3’ UTR (blue), which could express a small downstream open reading frame encoding a protein of unknown function.

Here, we focus on one IRES RNA sequence that emerged in this search, due to the peculiar nature of its genomic location and predicted structure. This putative IRES is located in the annotated 3’ UTR of penguin megrivirus (also called megrivirus E; MeV-E),^52^ within the *Picornaviridae* family (Fig. 1A). Specifically, the IRES is preceded by the main coding region (ORF1) and followed by nearly 400 nucleotides and a poly(A) tail (Fig. 1B). To date, no other Type IV IRES has been found outside of the 5’ UTR. This unexpected location for a Type IV IRES presents the question: why would an IRES be *downstream* of the main ORF and its stop codon, in a region annotated as untranslated RNA?

We first considered the possibility that this was an artefact from sequencing errors. However, the predicted 3’ IRES exists in all deposited sequencing reads of MeV-E and is not a duplication or assembly error as it only shares 15% primary sequence similarity with another Type IV IRES located in the 5’ UTR of the same genome.^52^ Taken together, it seemed unlikely for this sequence to be a false positive or artifactual.

If the RNA structure we identified in MeV-E is functional, it must initiate within what is annotated as untranslated region of the viral mRNA. Strikingly, the location of the AUG start codon for this putative 3’ IRES matches a start codon for an ORF encoding a putative 80 amino acid long peptide that could be expressed from the MeV-E 3’ UTR. While MeV-E is the only virus in which we identified a 3’ IRES, a second ORF (ORF2) encoding a putative small peptide has been proposed in the 3’ region of other megriviruses.^53,54^ The potential for an additional coding region in related viruses motivated us to continue our analysis of this newly identified MeV-E 3’ IRES RNA structure that could translate a downstream ORF through internal initiation.

In addition to its surprising genomic location, the structure of the MeV-E 3’ IRES is also highly divergent compared to the HCV IRES. While the core region of the MeV-E 3’ RNA is consistent with other Type IV IRESs,^37,38^ the rest appear to be dissimilar or missing entirely. Specifically, domain II is potentially much smaller compared to that of HCV and domain III is drastically truncated.

### The megrivirus E genome contains two IRESs capable of cap-independent translation initiation

Given the unique and curious nature of the MeV-E 3’ IRES RNA, we used translationally competent rabbit reticulocyte lysate with various dual luciferase reporter RNAs to test whether it could internally initiate translation in a cap-independent manner (Fig. 2A). These reporters were derived from the pSGDLuc system,^55^ which uses peptide-bond-skipping 2A sequences;^56^ this ensures that any translated sequences derived from control or viral RNA inserted between the two luciferase genes are liberated from the mature luciferase proteins and do not interfere with their luminescence signal. Additionally, testing RNAs prepared *in vitro* for translation initiation activity outside the context of a cellular environment avoids potential false positive signals that can result from cryptic promoters or splice sites.^57,58^

**FIGURE 2.**
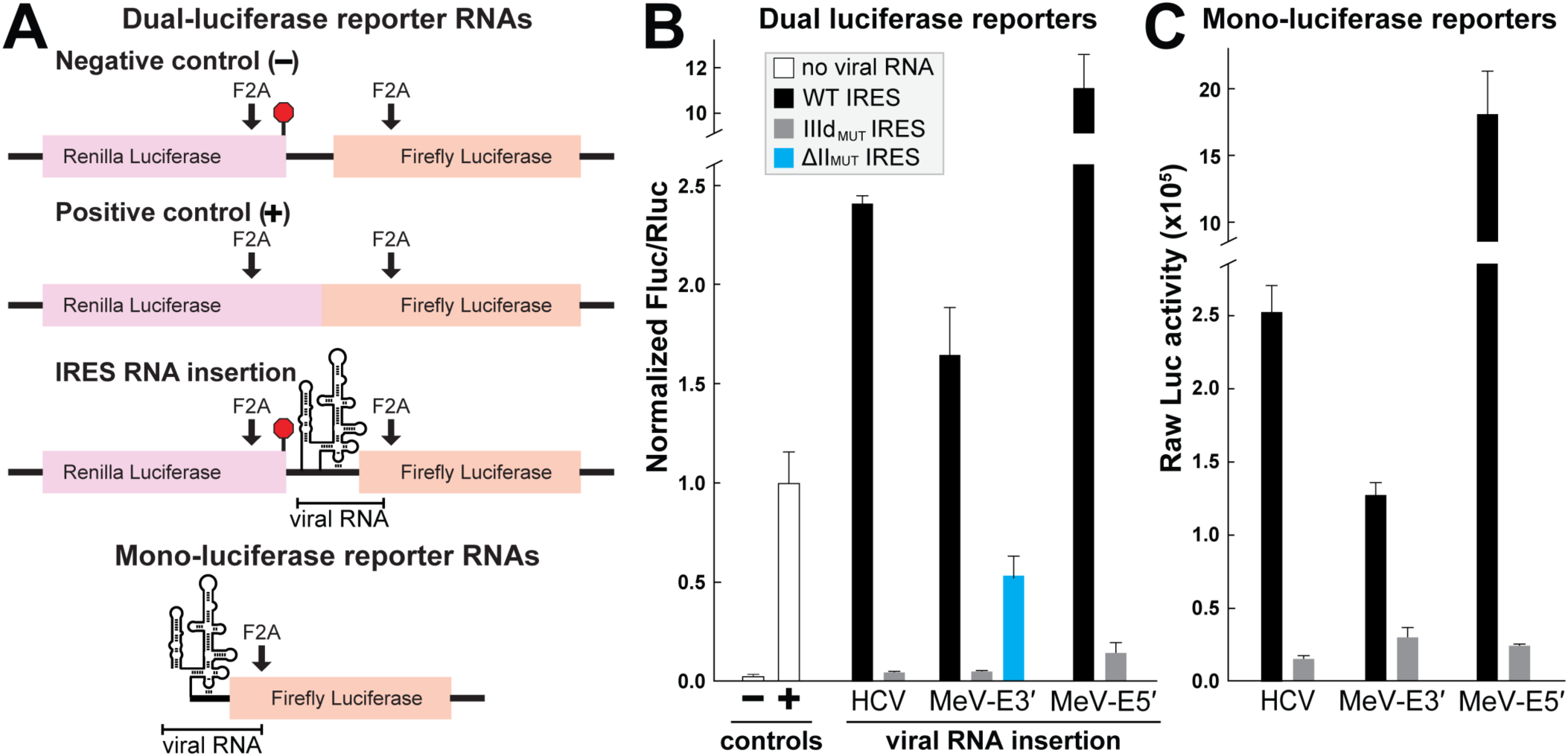
Both MeV-E IRESs are functional in promoting cap-independent translation initiation. (A) Design of control and experimental (IRES) dual- and mono-luciferase reporter RNAs. F2A sequences induce peptide bond-skipping events during translation. (B) Translation assays performed in rabbit reticulocyte lysate containing dual luciferase reporters carrying either no viral RNA (controls) or WT or mutant IRESs from HCV or MeV-E (experimental). The ratio of luminescence from the downstream (Fluc) to upstream (Rluc) proteins was normalized to the positive control. Error bars represent standard deviation. The locations of the IIId_MUT_ and ΔII_MUT_ truncation are displayed in Figure S1. (C) Translation assays performed in rabbit reticulocyte lysate containing firefly luciferase reporters carrying WT or mutant IRESs from HCV or MeV-E. Unnormalized luminescence values are plotted and error bars represent one standard deviation from the mean.

The MeV-E 3’ IRES construct tested contained MeV-E genomic sequence starting immediately after the ORF1 stop codon, through the predicted ORF2 start codon, ending 15 codons into ORF2 to ensure the entire putative IRES element was present. The normalized luminescence ratios measured for the wildtype (WT) HCV, MeV-E 5’, and MeV-E 3’ IRESs reflect their ability to directly initiate translation independent of the 5’ end of the reporter mRNA (Fig. 2B).

Three conserved guanosines in the loop of domain IIId are necessary for Type IV IRES function due to their role in 40S binding. Specifically, these guanosines base pair with cytidines in the apical loop of helix 26 (h26), also known as expansion segment 7, of the 18S rRNA.^20,21,31,39^ Consistent with previous observations, mutation of all three guanosines in the IIId stem loop (IIId_MUT_) ablates translation activity for the HCV, MeV-E 5’, and MeV-E 3’ IRESs (Fig. 2A, S1).

Due to its unusual genomic location, we also tested if the MeV-E 3’ IRES can initiate translation when placed in any position within the mRNA, not only an intergenic region adjacent to an upstream stop codon. When placed in the 5’ UTR of a mono-cistronic reporter RNA (Fig. 2A), the WT HCV, MeV-E 5’, and MeV-E 3’ IRESs all demonstrate translation activity well above their respective IIId_MUT_ counterparts. Taken together, the translation assay data led us to conclude that both the MeV-E 5’ and 3’ IRES RNAs are functional members of the Type IV IRES class.

While viruses in *Dicistroviridae* contain IRESs in both their 5’ UTR and intergenic region, they are not similar in structure.^4^ In contrast, the MeV-E genome is unique in that it contains two distinct IRES RNAs that both belong to the same type of IRES. The translation efficiency of the MeV-E 3’ IRES measured using lysate is lower than that of the HCV IRES and substantially lower than the MeV-E 5’ IRES for both mono- and dual-luciferase reporters RNAs. We discuss possible functional implications of disparate translation activity levels between the two MeV-E IRESs in a later section. After demonstrating that the MeV-E 3’ IRES is functional, we performed a variety of experiments interrogating its divergent structure compared to most Type IV IRESs to determine the mechanism underlying its relatively lower translation efficiency.

### Unusual architecture of the megrivirus E 3’ IRES RNA supported by structure probing

The initial secondary structure model for the MeV-E 3’ IRES, as determined by the output alignment from the homology search, predicts that several domains are missing compared to the HCV IRES (Fig. 3A,B). We used an *in vitro* chemical probing method^59^ to investigate whether this predicted secondary structure is correct. Chemical probing reactivities of the MeV-E 3’ IRES core region are consistent with the predicted secondary structure model including the two pseudoknots (Fig. 3C). Certain loop nucleotides within IIId and IIIe have high reactivity, leaving these sites available to form intermolecular base pairs with nucleotides in the loop of 18S rRNA h26, consistent with the HCV IRES in its ribosome-bound form.^31,32^

**FIGURE 3.**
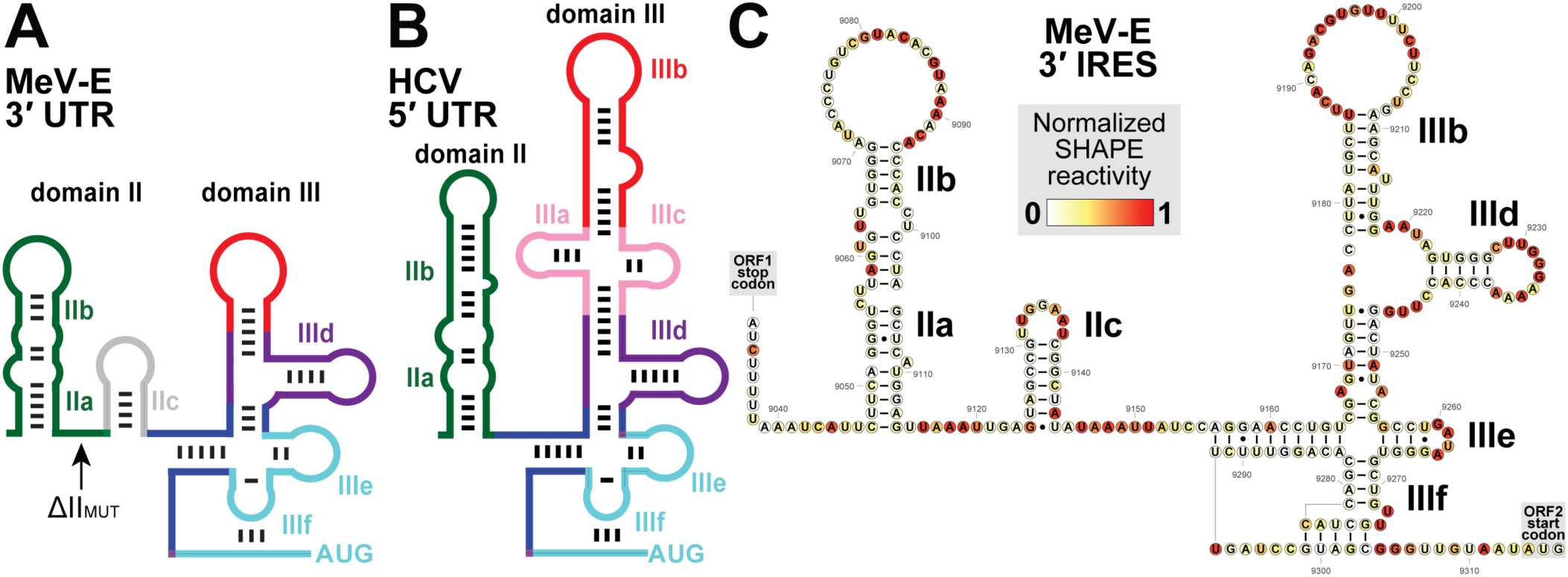
Smaller size and divergent domain architecture of the MeV-E 3’ IRES supported by chemical probing data. (A,B) Cartoon depictions of the secondary structure proposed for the MeV-E 3’ IRES (panel A) and the known structure of the HCV IRES (panel B). (C) Normalized reactivity at each nucleotide position to the chemical probing reagent 1-methyl-7-nitroisatoic anhydride (1M7) mapped onto a secondary structure model of the MeV-E 3’ IRES RNA.

Outside the core region, the MeV-E 3’ IRES structure substantially diverges from other Type IV IRESs. In lieu of domains IIIa, IIIb, and IIIc there is only one stem loop in the region extending past domain IIId. Chemical probing data are consistent with this secondary structure model in that base-paired nucleotides display very low reactivity while most of the 22 loop nucleotides display high reactivity. While we refer to this element as domain IIIb due to its location in the secondary structure, it may or may not serve the same functional role and/or occupy the same space as the HCV IRES domain IIIb in its ribosome-bound form.

The output alignment from the bioinformatic homology search predicted a very small stem loop corresponding to domain II. To test whether this small structure fully encompasses domain II and to define the minimal construct corresponding to the functional MeV-E 3’ IRES, we made a truncation to the RNA (ΔII_MUT_) (Fig. S1). This truncation results in a significant loss of translation activity compared to the WT construct (Fig. 2C), which contains an additional 80 nucleotides of sequence also present in our chemical probing RNA construct. Since this upstream sequence is necessary for full IRES activity, we favor a secondary structure model in which a larger stem loop, which we term domains IIa and IIb is located further upstream with an intervening “IIc” stem loop before domain III (Fig. 3, S1, S2). The secondary structure model of domain II was determined using the RNAfold program which took chemical probing reactivities into account for the prediction. Again, these stem loops may or may not serve the same functional role and/or occupy the same space as HCV IRES domain II. Notably, deletion of domain II of the HCV IRES is more deleterious to its translation activity compared to our ΔII_MUT_ MeV-E 3’ IRES.^36^

### Structure of the megrivirus E 3’ IRES RNA bound to the ribosome resolved by cryoEM

To better understand how the unusually compact MeV-E 3’ IRES folds and interacts with translation machinery, we used cryo-electron microscopy (cryoEM) to visualize an IRES-80S ribosomal complex assembled from purified components. We collected 11,782 micrographs and after many rounds of 2D and 3D classification narrowed down to 62,736 high-quality particles of the IRES-bound ribosome complex (Fig. S3). We did not observe any map features corresponding to the IRES interacting with the 60S portion of the ribosome, consistent with previous structures of the HCV IRES-80S complex,^32^ leading us to focus on the 40S subunit (Fig. S3). The resulting refined maps have features in a region of the 40S subunit adjacent to the mRNA exit channel beyond what is accounted for by the ribosome itself and we attribute this to the MeV-E 3’ IRES RNA (Fig. S4A). We fit an existing structure of the ribosome-bound HCV IRES^31^ into our experimental map to evaluate the extent of its similarities with the smaller MeV-E 3’ IRES (Fig. 4A).

**FIGURE 4.**
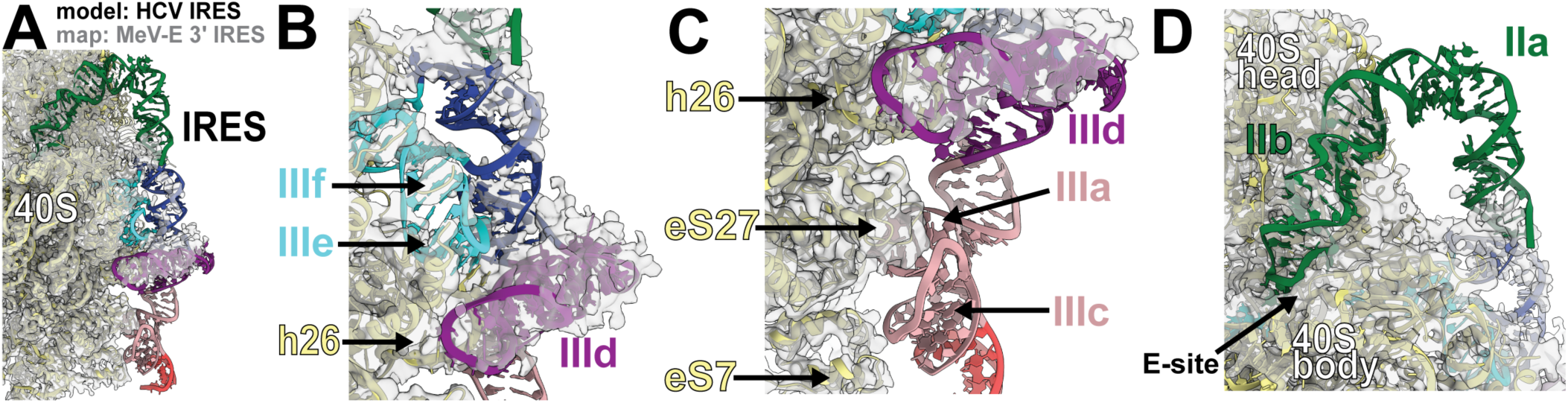
CryoEM reconstruction of the ribosome-bound MeV-E 3’ IRES reflects its smaller size and reduced binding capacity compared to the HCV IRES. (A) CryoEM map of the MeV-E 3’ IRES-ribosome complex fit with a previously determined structure of the 40S-HCV IRES complex (PDB: 5A2Q)^31^ to demonstrate the absence of density in specific regions. Domains of the HCV IRES are labeled and color coded consistent with Figure 3B. (B,C,D) Locally refined cryoEM map of the 40S-MeV-E 3’ IRES complex fit with the HCV IRES, zoomed in on the core region (panel B), domain III (panel C), and domain II (panel D). Ribosomal proteins (eS27, eS7) and regions of the 18S rRNA (helix 26; h26) are labeled in yellow.

Map features corresponding to the core region that is highly conserved in all Type IV IRESs are readily visible in our maps (Fig. 4B).^37,38^ This was expected as the secondary structure and many nucleotide identities in this region of MeV-E 3’ IRES are the same as the HCV IRES. Interactions between the loops of domains IIId and IIIe with helix 26 of the 18S rRNA (also known as ES7) of both IRESs appear largely similar (Fig. 4C). Strikingly, the map drops off abruptly after domain IIId (Fig. 4C). Overall, we see no map features corresponding to the apical portion of domain III of the MeV-E 3’ IRES. Taken together with our alignments and chemical probing data, our cryoEM maps provide additional evidence that structures equivalent to the HCV IIIa and IIIc domains are absent from the MeV-E 3’ IRES.

We were unable to resolve the majority of domain II, which is the other portion of the MeV-E 3’ IRES that is divergent compared to most Type IV IRESs. The map quality falls off before the base of this stem, which forms a coaxial stack on the base of domain III for the HCV IRES (Fig. 4D). There are minimal features in the E-site beyond what is accounted for by the ribosome (Fig. 4D), which is where the tip of domain II sits in the HCV IRES-40S structure.^22,26,33^ Thus, we did not assign any map features to specific nucleotides preceding domain III.

We are currently unable to make conclusions as to the nature of the divergent MeV-E domain II in terms of its 3D structure and ribosomal interactions. Our chemical probing data suggest that the linkers between domains IIab, IIc, and III are flexible (Fig. 3C). This flexibility likely allows domain II stem loops to occupy a diverse range of positions, and this heterogeneity leads to their absence from our cryoEM maps. The tip of domain IIb may occupy the E-site as it does for the HCV IRES, but this state was not well populated in our sample. Domain IIb of the MeV-E 3’ IRES does not contain a ‘loop E’ motif, which is present in many but not all Type IV IRESs (Fig. S1). A different complex, such as the MeV-E 3’ IRES bound only to the 40S subunit, might enable visualization of domain II by increasing occupancy of one state. Another possibility is that domain II of the MeV-E 3’ IRES does not occupy the E-site and instead performs other roles.

Given the apparent structural heterogeneity and limited local resolution in peripheral portions of the IRES (Fig. S3), we built a model only of the core region of the MeV-E 3’ IRES using inferred homology with the HCV IRES RNA (Fig. 5, S4). There are some map features adjacent to domain IIId unaccounted for by our model, which we expect is occupied by nucleotides connecting IIIb to the core region that we did not build (Fig. 5, dashed gray lines).

**FIGURE 5.**
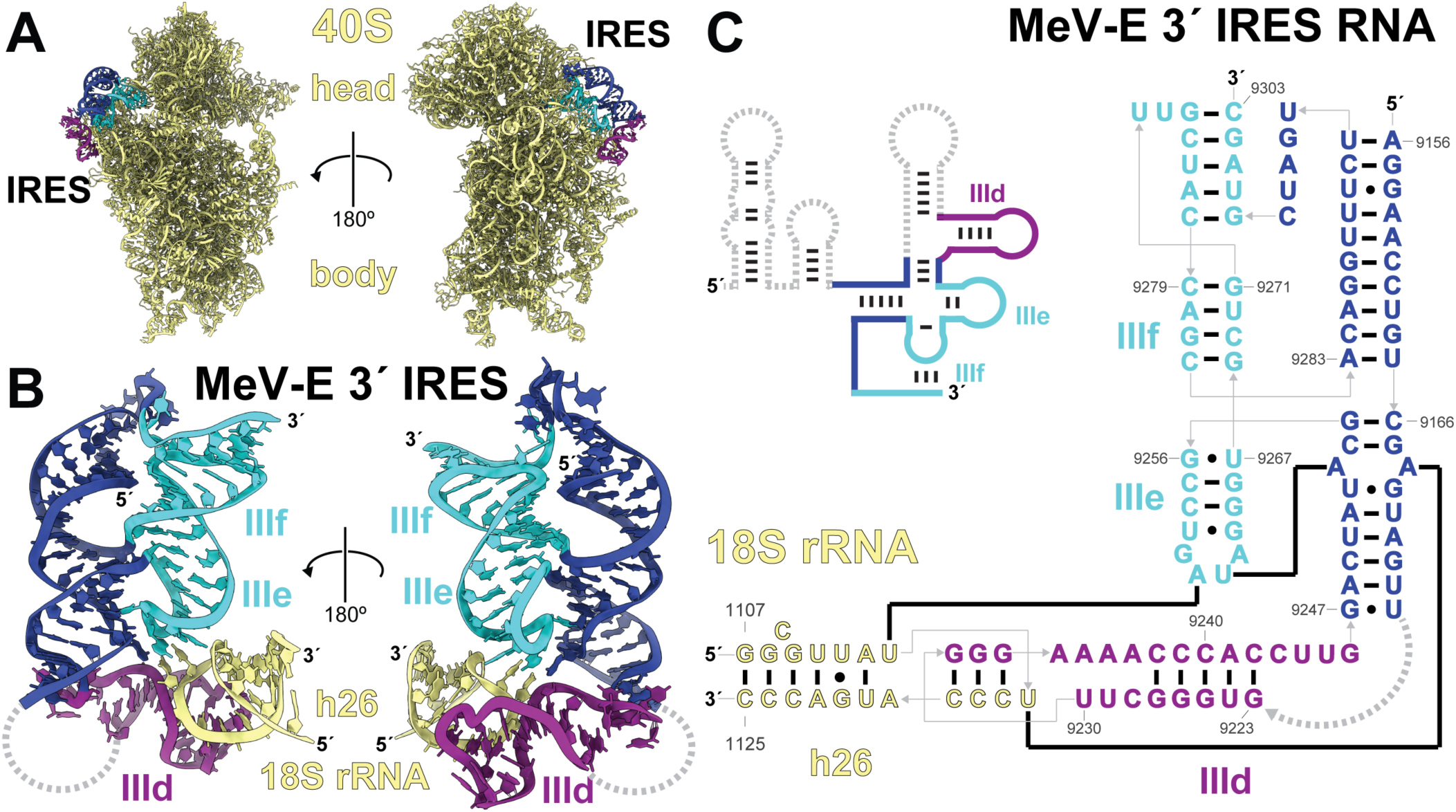
Model of the core region of the MeV-E 3’ IRES structure in its ribosome-bound state resolved by cryoEM. (A) Structural model of the portion of the MeV-E 3’ IRES RNA visualized by cryoEM, bound to the 40S ribosomal subunit (PDB: 9OGY). (B) Model of the 40S-bound MeV-E 3’ IRES RNA structure with its binding site on the apical portion of helix 26 (h26, also known as expansion segment 7) of the 18S rRNA (yellow). (C) Secondary structure depiction of the MeV-E 3’ IRES RNA bound to the apical portion of helix 26 of the 18S rRNA. Connectivity of the RNA is shown in gray and base pairing in black. The secondary structure cartoon depicts portions of the IRES RNA absent from the cryoEM reconstruction with dashed gray lines. Color coding of IRES domains is consistent with Figures 3 and 4.

### Impacts of missing domains on the MeV-E 3’ IRES initiation mechanism

Based on our Type IV IRES RNA structural alignment and chemical probing data, we anticipated that the MeV-E 3’ IRES would lack features corresponding to domains IIIa and IIIc in our cryoEM map. Indeed, there is no density in the regions these stem loops occupy (Fig. 4D). In the HCV IRES, the loops of these domains come together to bind ribosomal protein eS27.^26,31^ Therefore, the absence of equivalent structures in the MeV-E 3’ IRES RNA results in a loss of these interactions with the ribosome.

In the HCV IRES, the region of domain III that is truncated in the MeV-E 3’ IRES binds eIF3. Notably, map features corresponding to the apical portion of domain IIIb of the HCV IRES were not present in previous cryoEM reconstructions of 40S-IRES complexes,^26,31^ but were present in a 40S-IRES-eIF3 complex.^40^ We hypothesized that the lower activity of the MeV-E 3’ IRES in translation assays is due at least in part due to its inability to form some or all intermolecular interactions formed by the HCV IRES domain III. To investigate the impact of these missing protein-interaction domains on the mechanism of initiation by the MeV-E 3’ IRES, we performed binding experiments *in vitro* and pulldowns of IRES complexes from lysate.

We measured the affinity of the WT and IIId_MUT_ MeV-E 3’ IRES RNAs as well as WT HCV IRES for mammalian 40S subunits using purified components. As anticipated, ribosome binding by the MeV-E 3’ IRES is severely impaired compared to the HCV IRES, which has low nanomolar affinity for 40S subunits (Fig. S5). Due to experimental constraints, we could not test a high enough 40S concentration to saturate binding of either the WT or IIId_MUT_ MeV-E 3’ IRES but estimate their K_D_ values to be >500nM and >1 μM, respectively. This poorer affinity for the 40S subunit is consistent with that measured for a domain IIIabc truncation mutant of the HCV IRES.^21^

To evaluate the role of eIF3 in MeV-E 3’ IRES-mediated initiation, we incubated biotinylated IRES RNAs in lysate, pulled down assembled complexes by streptavidin affinity, and blotted for the presence of components of the eIF3 complex. The WT MeV-E 3’ IRES pulls down the 40S subunit along with multiple eIF3 proteins in a manner consistent with the HCV IRES (Fig. S6). A mutant RNA construct with a truncation of the apical portion of the MeV-E 3’ IRES domain III (ΔIIIb_MUT_; see Fig. S1) is also able to pull down the 40S subunit but loses association with components of the eIF3 complex (Fig. S6). These data implicate the MeV-E 3’ IRES domain IIIb as being functionally analogous to the IIIabc domain of the HCV IRES despite a substantial reduction in size, at least in terms of involvement in eIF3 biding. Every protein in the eIF3 complex that we blotted against was present in the MeV-E 3’ IRES preinitiation complex, but the molecular details of this interaction remain unclear. Additional biochemical or structural studies are needed to determine which residues within eIF3 proteins form direct interactions with nucleotides within the MeV-E 3’ IRES RNA. Perhaps eIF3a and eIF3c are directly bound and the core of the eIF3 complex is displaced from its canonical binding site, as with HCV IRES (Fig. 6A,B). Interestingly, the lack of interaction between the MeV-E 3’ IRES RNA and eS27 frees up the binding site for eIF3c on the 40S subunit, leading to additional possibilities for intermolecular interactions and the overall architecture of this IRES-40S-eIF3 preinitiation complex (Fig. 6C).

**FIGURE 6.**
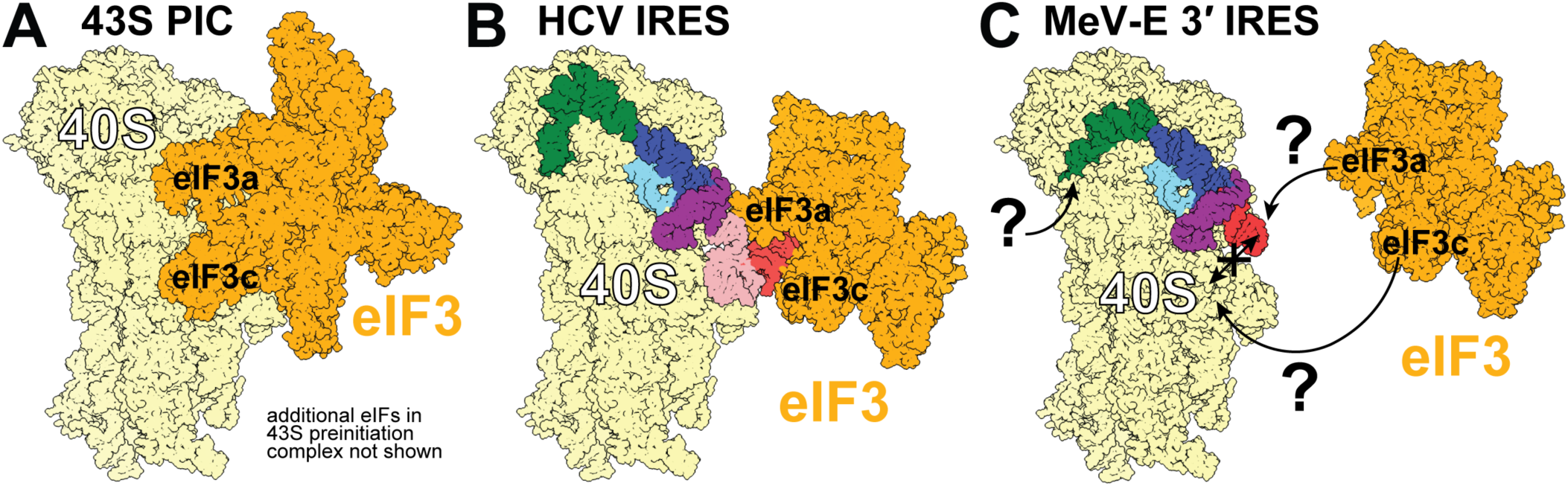
Canonical and non-canonical IRES-based modes of initiation using eIF3. (A) Canonical binding site of eIF3 on the 40S subunit (PDB: 5A5T)^81^ (B) Binding of eIF3 by the HCV IRES positions it away from its canonical binding site on the 40S subunit (PDBs: 5A2Q, 3J8B, 4C4Q; EMD-2451).^31,40,88^ (C) Speculative model of involvement of eIF3 during initiation by the MeV-E 3’ IRES, questioning whether eIF3 can access a portion of its canonical binding site on the 40S subunit or its interactions are mediated through the divergent domain IIIb (red) of the IRES (PDBs: **9OGY**, 3J8B).^88^ The structure and positioning of domain II (green) and IIIb (red) are hypothetical, modeled consistently with the HCV IRES.

Based on these results, we speculate that the pared down nature of domain III of the MeV-E 3’ IRES causes a loss of interactions with eS27 but not with the eIF3 complex. This weakens the affinity of the MeV-E 3’ IRES for the 40S subunit, affecting the step of initial ribosome recruitment and lowering the translation efficiency of this IRES compared to both HCV and the MeV-E 5’ IRES. Other aspects of the MeV-E 3’ IRES initiation remain mysterious. For example, the structure and functional role of its divergent domain II remain unclear. Additional studies are needed to tease apart whether any mechanistic differences involving domain II contribute to its lower translation activity.

### Structure, function, and conservation of a 3’-encoded peptide in megriviruses

After confirming that the MeV-E 3’ IRES is an authentic and functional member of the Type IV IRES class of RNAs, we examined the nature of the protein whose translation it controls. After analyzing the 3’ UTR sequence, we reannotated the region of the genome following ORF1 as a 283-nucleotide intergenic region, which includes the IRES RNA sequence, an 80 amino acid long coding region termed ORF2, and a 146-nucleotide 3’ UTR (Fig. 7A). We next used various bioinformatic tools to evaluate the characteristics and structure of the peptide sequence encoded by ORF2. One particularly striking feature is a strongly predicted alpha-helical transmembrane domain and topology that classifies it as a type I single pass transmembrane protein with the C-terminus in the cytoplasm (Fig. 7B). When mapped onto an independent protein folding prediction, residues that would span the membrane align to an alpha helical region (Fig. 7C).

**FIGURE 7.**
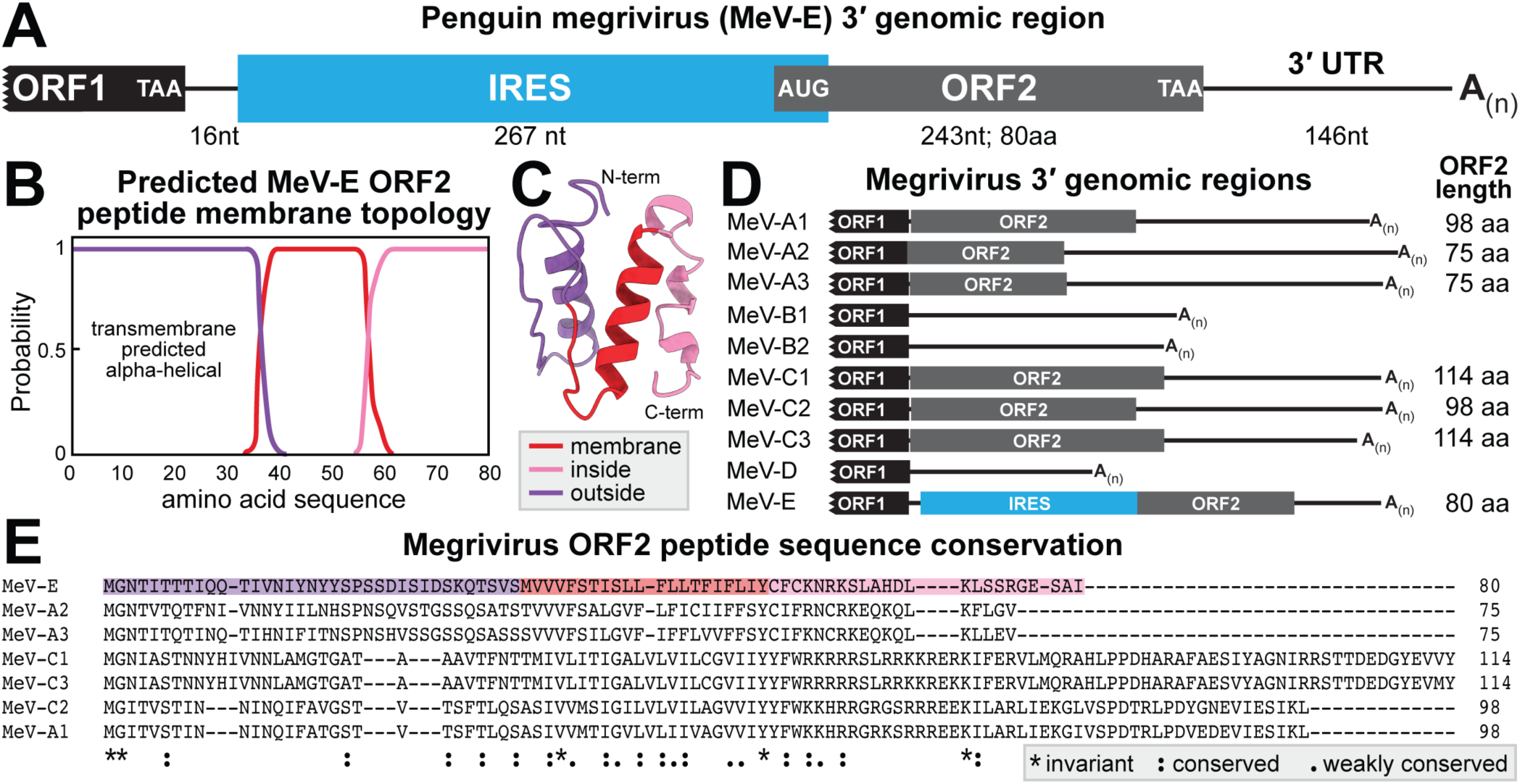
A second open reading frame conserved in some megriviruses encodes a small transmembrane protein. (A) Cartoon depiction of the portion of the MeV-E genome previously annotated as the 3’ UTR, which contains an IRES and small downstream open reading frame (ORF2). (B) Predicted transmembrane character and topology of the 80 amino acid protein encoded by ORF2 of MeV-E. (C) Predicted structure of the 80 amino acid protein encoded by ORF2 of MeV-E with color-coded membrane topology mapped to the structure. (D) Cartoon depiction of the annotated 3’ UTR of every virus within the Megrivirus genus, some of which contain a proposed second open reading frame. (E) Alignment of all proposed megrivirus ORF 2-encoded protein sequences.

As mentioned above, it was previously proposed that the genomes of both chicken megrivirus (MeV-C1) and turkey hepatitis virus (MeV-A1) contain a downstream ORF2, which would encode proteins 114 and 98 amino acids in length, respectively (Fig. 7D).^53,54^ While noting the coding potential in these 3’ regions, previous publications stated that the function of this protein as well as how is it translated are unknown.^53,54^ In contrast, the presence of the IRES in MeV-E led to our discovery of its ORF2.

We evaluated the length and protein-coding potential of the region 3’ of the ORF1 stop codon, previously annotated as 3’ UTRs, in all megriviruses (Fig. 7D). Previous publications stated that duck megrivirus MeV-A2, goose megrivirus (MeV-A3), pigeon mesivirus 1 and 2 (MeV-B1, MeV-B2), and harrier megrivirus (MeV-D) do not contain an additional ORF in this 3’ region.^60–63^ Our analysis supports that absence of an ORF in MeV-B2, -B3, and -D, which all have shorter 3’ regions. However, we identified a putative 75 amino acid ORF2 in both MeV-A2 and -A3 (Fig. 7D). For MeV-A2, this putative ORF2 was likely missed because its start codon overlaps and shares nucleotides with the ORF1 stop codon.

We aligned all likely megrivirus ORF2 peptide sequences and found regions of high conservation, including the alpha helical transmembrane domain predicted for the MeV-E protein (Fig. 7E). The N-terminal regions display some conservation and are similar in length while the C-terminal regions, predicted to be cytoplasmic, are less conserved and quite variable in length.

Many other viruses, including influenza- and coronaviruses, encode short peptides with a single pass transmembrane domain that multimerize to form channels through membranes; this type of protein is called a viroporin.^64,65^ Members of the *Caliciviridae* family contain a small downstream ORF encoding the minor capsid protein, which decorates the surface of the viral capsid surface and enables delivery of the viral genome through the endosomal membrane into the cytoplasm.^66^ Notably, some megriviruses have the stop codon of ORF1 overlapping or proximal to the start codon of ORF2. This arrangement of ORF1 and ORF2 is similar to that of influenza B virus and many caliciviruses. These viruses use an RNA structure to induce translation termination-reinitiation^67^ to express ORF2, which is translated at lower levels compared to the main ORF.^49,68,69^ Reinitiation could also be the mechanism employed by MeV-A and -C, which contain ORF2 but lack an IRES, although other types of noncanonical translation cannot be ruled out. While reinitiation differs from IRES-driven internal initiation in its mechanism,^49,70^ these two types of RNA structure-mediated translation can accomplish the same functional outcome in expressing downstream ORFs with lower levels of ORF2 compared to ORF1.

There are likely many unannotated viral open reading frames waiting to be discovered that are hidden due to their reliance on alternative translation events. In our case, an exploration of RNA structural diversity led to the discovery of a new viral protein whose role during infection and impact on host cell specificity can now be explored.

### Tuning translation through RNA structural modularity

The power of structure-guided RNA homology searches is exemplified by the advances we report in this manuscript. This bioinformatic strategy enabled identification of the first Type IV IRES located outside of a 5’ UTR, which was unlikely to have been found by other means. The pared down nature of the MeV-E 3’ IRES RNA structure has several broader implications for translation initiation mechanism and regulation.

We measured a ∼10-fold difference in translation activity between the two IRESs within the MeV-E genome using reporter RNAs in lysate. In the context of viral infection, this would cause lower-level expression of the ORF2 peptide compared to the rest of the viral genome encoded by ORF1, resulting in a 10:1 stoichiometry of protein products. While a mammalian lysate may not faithfully represent the exact expression levels during viral infection within a bird cell, the translation machinery these IRESs interact with is highly conserved. For example, the sequence of 18S rRNA helix 26, which serves as a key site of interaction through intermolecular base pairing with the IRES, is identical throughout vertebrates. Regardless of the exact ratio of proteins, we expect that the pattern of reduced translation at ORF2 compared to ORF1 is likely to be true in infection. We speculate that this disparity is purposeful for the virus, and the IRES structures evolved to create a beneficial differential in protein levels. This “dual-IRES” arrangement is another strategy in the viral toolbox to regulate protein expression through RNA structure.

Entire domains necessary for efficient translation by the HCV IRES have been trimmed down or otherwise compensated for by the MeV-E 3’ IRES, calling into question whether all Type IV IRESs use the same mechanism to initiate translation. While studying one sequence as a model system can be hugely beneficial, ignoring heterogeneity prevents us from tapping into the full potential and variety embodied by a class of RNA structures. Our findings warrant more detailed experiments linking translation efficiency with 3D structure, intermolecular interactions, and preinitiation complex composition of many structurally divergent members of the Type IV IRES class. This line of research could open numerous possibilities with intriguing biotechnology implications. One particularly desirable feature of the MeV-E 3’ IRES is its smaller size compared to the HCV IRES. Designing a minimal Type IV IRES would be valuable for use in reporters or mRNA therapeutics when the number of nucleotides is at a premium.

Defining the function and efficiency of different versions of each domain could enable rational design of IRESs with desired translation levels in a range of cell types and under stress conditions. We envision this could create an assortment of “plug and play” Type IV IRESs with tunable expression through various combinations of RNA structural modules. Teasing apart RNA structure-functions relationships will continue to enable fundamental biological discoveries as well as the development of molecular tools with beneficial biotechnology applications.

## Resource availability

### Lead Contact

Requests for further information and resources should be directed to and will be fulfilled by the lead contact Jeffrey Kieft.

### Material availability

Plasmids generated in this study will be made available on request.

### Data and code availability

- The Cryo-EM maps for Megrivirus E 3’ internal ribosome entry site (IRES) RNA core region bound to rabbit ribosome have been deposited in the Electron Microscopy Data Bank (EMDB) under accession code EMD-70479. Coordinates for the resultant model have been deposited in the Protein Data Bank (PDB) under accession number 9OGY.
- This study does not report any custom code.
- Any additional information required to reanalyze the data reported in this paper is available from the lead contact upon request.

## Acknowledgements

We thank current and former members of the Kieft laboratory for helpful discussions. We thank Dr. Daniel DiMaio for advice and insight on viral miniproteins. The pSGDlucV3.0 was a gift from John Atkins (Addgene plasmid # 119760). This work was supported by National Institutes of Health/National Institute of General Medical Sciences (NIH/NIGMS) grant R35GM118070 and a McKnight Foundation Award to

JSK. A portion of this research was supported by NIH grant R24GM154185 and performed at the Pacific Northwest Center for Cryo-EM (PNCC) with assistance from Vamsee Rayaprolu. MES was a Jane Coffin Childs Postdoctoral Fellow. KES was supported by NIH grants T32GM136444 and F31AI186389-01.

## Author contributions

MES: Conceptualization, Data curation, Formal Analysis, Funding acquisition, Investigation, Methodology, Visualization, Writing – original draft, Writing – review & editing. KES: Investigation, Methodology, Visualization, Writing – review & editing. JSK: Funding acquisition, Supervision, Visualization, Writing – review & editing.

## METHOD DETAILS

### Bioinformatic identification of Type IV IRESs

Existing alignments of Type IV IRESs were obtained from the Rfam database,^71^ originally called the Hepatitis C Virus IRES (Rfam ID: 00061) and Pestivirus IRES (Rfam ID: 00209) classes, and their secondary structure models were merged. Four additional IRES sequences whose secondary structures had been experimentally validated (porcine kobuvirus, GB virus B, simian picornavirus 1, and porcine teschovirus) were added to the alignment. This seed alignment was used to query a database of consisting of all +ssRNA virus sequences deposited in the National Center for Biotechnology Information (NCBI, retrieved 01/22/2019) using the program Infernal.^42^ This homology search returned additional examples of the Type IV IRES motif, including the MeV-E 3’ IRES described in this study. Additional details about this bioinformatic search are described elsewhere.^51^

### RNA preparation for dual luciferase assays

Dual luciferase constructs containing intervening viral RNAs of interest were made using the pSGDLucV3.0 vector (Addgene 119760). Insertion sequences were ordered as double-stranded DNA gene fragments (IDT) then amplified via PCR. Vector and inserts were prepared by digesting with BglII (NEB) in NEBuffer r3.1 (NEB) at 37 °C for 2 hours followed by digestion with PspXI (NEB) in rCutsmart™ Buffer (NEB) and incubated at 37 °C for an additional 2 hours. The vector was purified via agarose gel purification while inserts were purified directly from the restriction digest reaction, both using the Wizard SV Gel and PCR Clean-up System (Promega). Inserts were incorporated into the vector via ligation reaction consisting of inserts and vector in a 5:1 molar ratio, T4 ligase (NEB), and T4 ligation buffer (NEB). The reaction was incubated at 16 °C overnight then transformed into DH5⍺ E. coli and screened for successful ligation via colony PCR. Sequences were verified via whole plasmid sequencing prior to use (Plasmidsaurus). The positive control (in-frame readthrough) and dIII_MUT_-containing plasmids were constructed using the Q5 site-directed mutagenesis kit (NEB) according to manufacturer protocol and verified via whole plasmid sequencing. Double-stranded DNAs approximately 5 kb (dual-luciferase) or 3 kb (mono-luciferase) in length were produced via PCR as a template and their size verified by 1.5% agarose gel electrophoresis. These PCR products were then used as templates for *in vitro* transcription with the MEGAscript™ T7 transcription kit (Invitrogen). RNA was purified using the Monarch RNA clean-up kit (NEB) and RNA length and quality was verified via 1.5% agarose gel.

### Translation assays using dual luciferase reporters in lysate

Each reaction contained 500 ng of RNA, which was initially heat re-folded by incubating in 33mM HEPES (pH 7.5) at 95 °C for 1 minute then cooling to room temperature for 5 minutes at which time MgCl_2_ was added to a final concentration of 10 mM. The final 50 µL translation reaction included the 5 µL RNA mixture described above and components from the Rabbit Reticulocyte Lysate system (Promega): 35 µL of rabbit reticulocyte lysate, amino acid mixture minus cysteine (final concentration 16.7 µM), amino acid mixture minus methionine (final concentration 16.7 µM), amino acid mixture minus leucine (final concentration 16.7 µM), and KOAc (final concentration 150 mM). Reactions were incubated at 30 °C for 4 hours. After incubation, reactions were diluted 4X with Passive Lysis Buffer (Promega) and the reaction was split into two technical replicates, each with a final volume of 100 µL. Luciferase activity was measured with a GloMax®-Multi Detection system using the Dual-Glo® Luciferase Assay System (Promega). Results were analyzed using Excel (Microsoft) software with measurements from at least three independent experiments, using the average of the two technical replicates from each trial. These measured values were used to calculate the ratio of Firefly (downstream) to Renilla (upstream) translation (Fluc/Rluc), then these ratios were normalized with the positive control (in-frame readthrough) set to 1. For mono-luciferase constructs, raw luminescence values corresponding to signal from the Firefly luciferase gene are reported as the average and standard deviation without any normalization. Data were plotted and labeled using Adobe Illustrator.

### RNA preparation for chemical probing, filter binding, immunoprecipitation, and cryoEM

Template DNA was acquired as gBlock DNA fragments (IDT) containing an upstream T7 promoter and amplified in PCR reactions under the following conditions: 100 ng plasmid DNA, 0.5 µM forward and reverse DNA primers (see Table S1), 500 µM dNTPs, 25 mM TAPS-HCl (pH 9.3), 50 mM KCl, 2 mM MgCl_2_, 1 mM β-mercaptoethanol, and Phusion DNA polymerase (New England BioLabs). Successful amplification of dsDNA was confirmed by 1.5% agarose gel electrophoresis. Transcriptions were performed in 5 mL volume using 1.6 mL of PCR product as template dsDNA, 30 mM Tris pH 8.0, 60 mM MgCl_2_, 8 mM each NTP, 10 mM DTT, 0.1% spermidine, 0.1% Triton X-100, and T7 RNA polymerase. Transcription reactions were incubated at 37°C overnight, then mixed with equal volume of a 2X loading dye (0.09 M tris, 0.09 M borate, 10 mM EDTA (pH 8.0), 18 M urea, 20% sucrose, 0.05% bromophenol blue, 0.05% xylene cyanol) and purified via denaturing 8% polyacrylamide gel electrophoresis (PAGE). RNA bands were detected via UV shadowing and subsequently excised. These bands were then extracted by crush soaking in diethylpyrocarbonate-treated (DEPC) milli-Q water at 4 °C overnight. The RNA-containing supernatant was concentrated using spin concentrators (Amicon) with a 30 kDa cutoff to achieve the desired RNA concentration in DEPC-treated water.

### *In vitro* chemical probing of RNAs using mutational profiling

Structure probing experiments using the chemical modifier 1-methyl-7-nitroisatoic anhydride (1M7) were performed as described in the published protocol describing the Selective 2’-hydroxyl acylation analyzed by primer extension and mutational profiling (SHAPE-MaP) method.^59^ Reactivities were the average of four replicates. Data were visualized using VARNA.^72^

### RNA biotinylation for immunoprecipitation

RNA prepared by *in vitro* transcription was subjected to ethanol precipitation then resuspended in an oxidation solution containing 100 mM sodium periodate and 100 mM NaOAc (pH 5.2) to a final RNA concentration of 1 µg/µL. Following incubation for 90 minutes at 23 °C, NaCl was added to 250 mM final concentration and reactions were placed on ice for 10 minutes. Reactions were spun down at 4°C for 5 min at 20,000 g and supernatant was retained. An equal volume of 100 mM BiotinXhydrazide (Lumiprobe) in DMSO was added, then biotinylated RNA was incubated at room temperature for 4 hours. Samples were then purified via Acid-Phenol:Choloroform (pH 4.5, Invitrogen) extraction, pelleted via ethanol precipitation, then resuspended in water to a final RNA concentration of 1 µg/µL.

### Immunoprecipitation and Western blotting

High-capacity streptavidin agarose resin (Thermo) was spun down at 500 g for 1 minute then storage buffer was removed. Beads were then washed three times using a buffer containing 20 mM HEPES-KOH (pH 7.6), 100 mM KOAc (pH 7.6), and 2 mM Mg(OAc)_2_ then resuspended with 1X sample volume of buffer. 120 µL of washed beads was added to 20 µL of heat refolded biotinylated RNA and incubated at 23 °C for 30 minutes with vigorous shaking to keep beads in suspension. After incubation, beads were spun down at 500 g for 1 minute, let stand for an additional 2 min then supernatant was removed.

Clarified lysate mixtures were prepared by spinning Hela cytoplasmic extract (Ipracell) at 12,000 g for 10 minutes at 4 °C, taking the supernatant and diluting to the following concentrations: 89% v/v lysate, 2 mM Guanosine 5′-[β,γ-imido]triphosphate (Sigma), 1X protease inhibitor cocktail (Roche). 120 µl of the lysate mixture was added to the prepared beads containing biotinylated RNA, incubating at 23 °C for 45 minutes with vigorous shaking. Beads were spun down at 500 g for 1 minute, let stand at 23 °C for 2 minutes, then supernatant was removed. Beads were washed 5X with 500 µL of a buffer containing 20 mM HEPES-KOH (pH 7.6), 200 mM KOAc (pH 7.6), 2 mM Mg(OAc)_2_, followed by one final wash with a buffer containing 20 mM HEPES-KOH (pH 7.6), 100 mM KOAc (pH 7.6), and 2 mM Mg(OAc)_2_. All wash buffer was removed, then 90 µL of loading buffer (4% w/v SDS, 250 mM Tris-HCl pH 6.8, 1.7 mM DTT, 30% v/v glycerol, 0.05% Bromophenol blue) was added to the beads.

Samples were heated to 95 °C for 5 minutes and mixed well before loading into 4-20% polyacrylamide gel (BioRad), which was run at 170 V for 50 minutes. Gels were transferred onto 0.45 µm Nitrocellulose membranes (Biorad) using the Trans-Blot® Turbo™ Transfer System at 25 V (1.0 Amp) for 30 minutes using a buffer containing 25 mM Tris-HCl (pH 7.6), 192 mM glycine, and 20% methanol. Membranes were blocked using 5% w/v milk in 20 mM Tris (pH 7.6) and 150 mM NaCl for 1 hour then washed twice with 20 mM Tris (pH 7.6), 150 mM NaCl, 0.1% w/v Tween 20. Primary antibody solutions were made with 3% w/v bovine serum albumin (Sigma) in 20 mM Tris (pH 7.6), 150 mM NaCl, 0.1% w/v Tween 20. Membranes were incubated overnight at 4°C with primary antibody mixtures (1:1000 dilutions) against: eIF3a (Invitrogen), eIF3b (Invitrogen), eIF3c (Proteintech), eIF3d (Proteintech), eIF3h (Proteintech), eIF3m (Proteintech), uS3 (Invitrogen).

After primary antibody solutions were removed, the membrane was washed 3X as described above. IRDye® 800CW Goat anti-Rabbit IgG Secondary Antibody (LI-COR) was diluted 1:20,000 in 3% bovine serum albumin w/v in 20 mM Tris (pH 7.6), 150 mM NaCl, 0.1% w/v Tween 20. Membranes were incubated with the secondary antibody mixture at 23 °C for 2 hours. After removing secondary antibody, the membrane was washed 3X, then blots were imaged on a LI-COR CLX Odessey. Figures containing scanned images were labeled and modified using Adobe Illustrator.

### Ribosome purification from lysate for cryoEM and filter binding

Nuclease-treated bulk rabbit reticulocyte lysate (Green Hectares) was spun down at 20,000 RPM for 15 min to remove debris, nuclei, and mitochondria.^73^ Clarified lysate was filtered with a 0.22 µm filter (Millipore). The supernatant was loaded on to a 30% sucrose cushion (20 mM Tris-HCl pH 7.5, 2 mM MgOAc2, 150 mM KCl, 30% w/v sucrose) and ultracentrifuged for 17.5 hours at 36,000 RPM (50.2 Ti rotor) at 4 °C to obtain a ribosomal pellet.^74^ The pellet was washed and resuspended in a buffer containing 20 mM Tris-HCl pH 7.5, 6 mM MgOAc_2_, 150 mM (80S purification) or 500mM (40S purification) KCl, 6.8% w/v sucrose, 1 mM DTT, 1 μL RNasin from Promega (Cat N2618).

To remove non-resuspended particles, the resuspended pellet was centrifuged again at 10,000 g for 10min at 4 °C and the supernatant was isolated. 15-30% sucrose gradients were prepared using a buffer containing 20 mM Tris-HCl pH 7.5, 2 mM MgOAc_2_, 150 mM (80S purification) or 500mM (40S purification) KCl, and either 15% or 30% w/v sucrose using the Gradient Master (Biocomp). Gradients were cooled to 4 °C before use. Pellet supernatant was loaded onto gradients and ultracentrifuged at 19,100 RPM for 17.5 hours (SW-28 rotor) at 4 °C. Gradients were fractionated using the Piston Gradient Fractionator and Fractionator Software v8.04 (Biocomp), monitoring for absorption at 260 nm.

Fractions corresponding to either pure 80S ribosomes or 40S subunits were separately collected and pooled. 80S ribosomes were concentrated to an A_260_ of 95 using an Ultra-15 centrifugal filter unit with a nominal molecular weight limit of 100 kDa (Amicon). This concentrated stock was subsequently diluted to 200 nM using buffer containing 20 mM Tris-HCl pH 7.5, 2 mM MgOAc_2_, 150 mM KCl and 20 μL aliquots were flash frozen in liquid nitrogen and stored at -80 °C. 40S subunits were concentrated by ultracentrifugation for 18 hours at 36,000 RPM (50.2 Ti rotor) at 4 °C and the ribosomal pellet was resuspended to an A_260_ of 360 in buffer containing 20 mM Tris-HCl pH 7.5, 2 mM MgOAc_2_, 150 mM KCl. Aliquots were flash frozen in liquid nitrogen and stored at -80 °C, then thawed and diluted to the desired concentration in the same buffer for filter binding experiments.

### Preparation of 5′-^32^P-labeled RNAs

RNAs prepared by *in vitro* transcription were dephosphorylated by combining 2 µL of 1 µM RNA, 2 µL 10X Antarctic Phosphatase buffer (NEB), and 14 µl of water. This mixture was heated to 95°C for 2 minutes then allowed to cool to room temperature. Then 2 µl of Antarctic Phosphatase (NEB) was added and the reaction was incubated at 37 °C for 1 hour followed by 2 minutes at 80 °C to heat inactivate the enzyme. Dephosphorylated RNA was radiolabeled by adding 10 µL of the previous reaction to 2.5 µL of 10X T4 Polynucleotide Kinase buffer (NEB), 1 µL T4 Polynucleotide Kinase (NEB), 1 µL ATP [γ-^32^P] (Revity), and 10.5 µL of water then incubating at 37 °C for 30 minutes then 65°C for 20 minutes. RNA was purified via denaturing 8% PAGE as described above and resuspended in water to 100 counts per minute.

### Filter binding

Each 50 µL filter binding reaction containing 2 µL radiolabeled RNA, 20 mM Tris-HCl (pH 7.6), 4 mM Mg(OAc)_2_, 300 mM KCl, 5% w/v sucrose, and 40S subunit concentrations ranging from 0 nM to 1 µM was incubated at 37°C for 15 minutes. Membranes were prepared by soaking in buffer containing 20 mM Tris-HCl (pH 7.6), 4 mM Mg(OAc)_2_, 150 mM KCl, and 5% w/v sucrose and assembled in the following order from bottom to top: Whatman® cellulose filter paper (Sigma), Amersham™ Hybond (Cytiva), 0.45 µm nitrocellulose membrane (Biorad), and Supor™ 0.45 µm membrane (Pall). The filter binding reactions were applied directly to the assembled membranes under vacuum followed by washing twice with the same buffer as above. Membranes were dried under vacuum, removed, and visualized on a Typhoon FLA 9500 Phosphoimager (GE).

Fraction bound values were calculated by quantifying the Hybond and nitrocellulose membranes. The apparent *K*_D_ values were determined by plotting the average and standard deviation of fraction bound from three separate replicates as a function of the logarithm of 40S subunit concentration. Data were fit using a symmetric sigmoidal dose response curve with the assumption of 1:1 binding (GraphPad Prism). For the MeV-E 3’ IRES RNAs, the fraction bound values are estimated since binding was not saturated at the highest concentration (1 µM) of 40S subunits.

### cryoEM sample preparation and image acquisition

MeV-E 3’ IRES RNA at 10 μM was heat denatured at 95 °C and allowed to cool slowly to ambient temperature, then 2 μL of the RNA solution was added to a 20 μL aliquot of 200 nM 80S and incubated at 37°C for 20 minutes. Using a FEI Vitrobot mark IV, 3 μl of the MeV-E 3’ IRES-80S solution was applied to grids (holey carbon grids C-flat 1.2/1.3 400 mesh VWR) that had undergone plasma cleaning for 6 seconds in a mixture of O2 and H2 by a Gatan Solarus 950 (Gatan, Inc). Grids were blotted for 4 seconds at a force of 0, then plunge frozen in liquid ethane.

Movies were collected using a Thermo Scientific 300 kV Krios G3i microscope equipped with a Gatan K3 Camera and Biocontinuum Energy Filter. A total of 11,782 micrographs were collected (4095 at 30° stage tilt to overcome preferred orientations) with a total dose of 50 e-/Å2. SerialEM was used to monitor data collection. Additional details can be found in Table S2.

### cryoEM image processing and map generation

Cryo-EM data were processed using cryoSPARC 4.0.^75^ An overview of the processing workflow can be found in Fig. S3. Movies were imported then processed (patch motion correction, patch CTF estimation) using default parameters. The 80S portion of an existing structure (PDB: 9C8K) was used to generate templates for particle picking. Particles were extracted from micrographs with a box size of 1024 x 1024 pixels, then downsampled 2X. Five initial rounds of 2D classification were performed to remove “junk” particles (e.g. incorrectly picked particles, ice contamination, aggregates). Particles from all good classes were used for ab initio 3D reconstruction requesting two classes followed by heterogeneous refinement resulting in one class with good 80S density and another demonstrating good 60S density but poor 40S density and those particles were removed. Four additional rounds of 2D classification were performed to remove particles belonging to classes with poor resolution and/or alignment.

All resulting good particle locations underwent local motion correction^76^ then the 60S signal was subtracted, and the masked 40S underwent non-uniform refinement. A mask was generated for the local region surrounding the IRES and including the decoding groove of the ribosome using UCSF Chimera.^77^ Focused 3D classification was performed with the IRES mask using varying numbers of classes. Ultimately two major classes were identified, one which contained a P-site tRNA and displayed poorer IRES density and these particles were excluded.

The remaining good 62,736 particles underwent non-uniform refinement, first with the 40S mask then subsequent local refinement with the IRES mask. Resolutions for each of these refinements (Fig. S3) were estimated using the gold standard Fourier shell correlation (GSFSC) of 0.143.

### Model building and refinement

A previous ribosome-bound HCV IRES structures (PDB: 5A2Q)^31^ was fit into the density of the map resulting from the non-uniform 40S mask refinement using ChimeraX.^78^ The IRES chain was split into helical element segments, which were mutated to reflect the sequence of the MeV-E 3’ IRES using COOT.^79^ These IRES segments were each individually refined then the backbone was reconnected including any linker nucleotides. This IRES chain was then recombined with a previous structure of the 40S subunit in its IRES-bound form (PDB: 7SYP)^33^ and further refined against the 40S map in Phenix.^80^

Modeling in Figure 6B was accomplished by fitting a structure of eIF3 (PDB: 5A5T)^81^ and the HCV IRES-40S structure (PDB: 5A2Q)^31^ into a map of a 40S-IRES-eIF3 complex (EMD-2451).^40^ Figure 6A shows only the 40S subunit and core octamer of the eIF3 complex from a previously solved preinitiation complex structure (PDB: 6ZMW).^82^ Figure 6C is the MeV-E 3’ IRES (PDB: 9OGY) with hypothetical structures of domains II and IIIb modeled and the positioning of eIF3 does not reflect a specific bound state. Figures of cryoEM maps and structural models were made using ChimeraX and labeled using Adobe Illustrator.

### Protein structure prediction

Sequences of putative ORF2 peptides were generated with NCBI ORF Finder (https://www.ncbi.nlm.nih.gov/orffinder/) using each annotated 3’ UTR sequence as the input. Accession numbers for the sequences used for each virus are listed in Table S3. Transmembrane character and topology was predicted using DeepTMHMM.^83^ Protein structure prediction was performed using I-TASSER^84–86^ and a figure of the resulting model was created using ChimeraX. Protein sequence alignments were created using Clustal Omega^87^ using default options for depicting conservation.

## SUPPLEMENTAL MATERIAL

Supplemental material is available for this article.

## Supplemental Information

**Figure S1.**
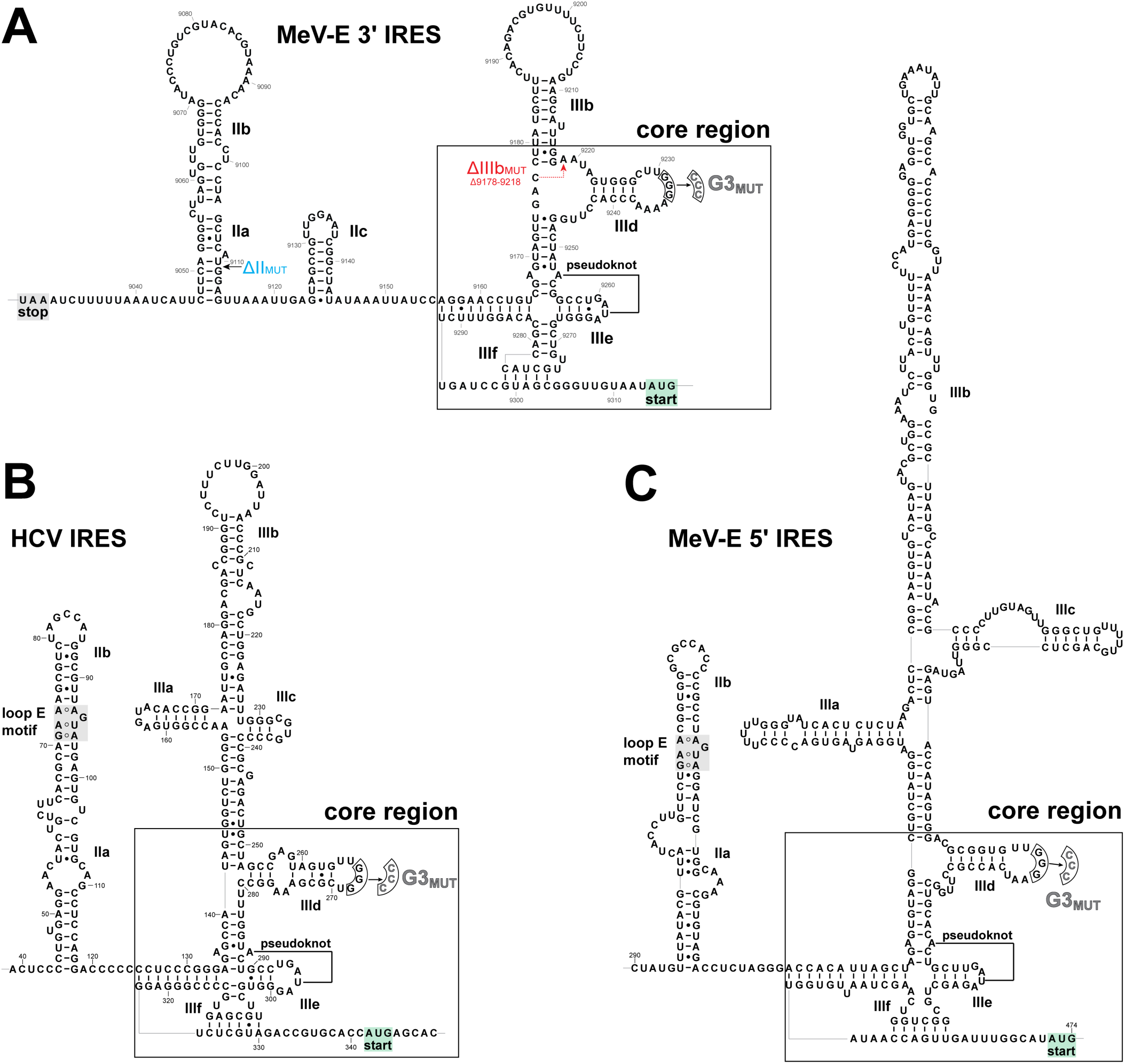
Secondary structure models of the HCV, MeV-E 5’, and MeV-E 3’ IRES RNAs. (A,B,C) Sequence and secondary structure models of the MeV-E 3’ (panel A), HCV (panel B), and MeV-E 5’ (panel C) IRESs displaying genomic coordinates and locations of mutations used in this study.

**Figure S2.**
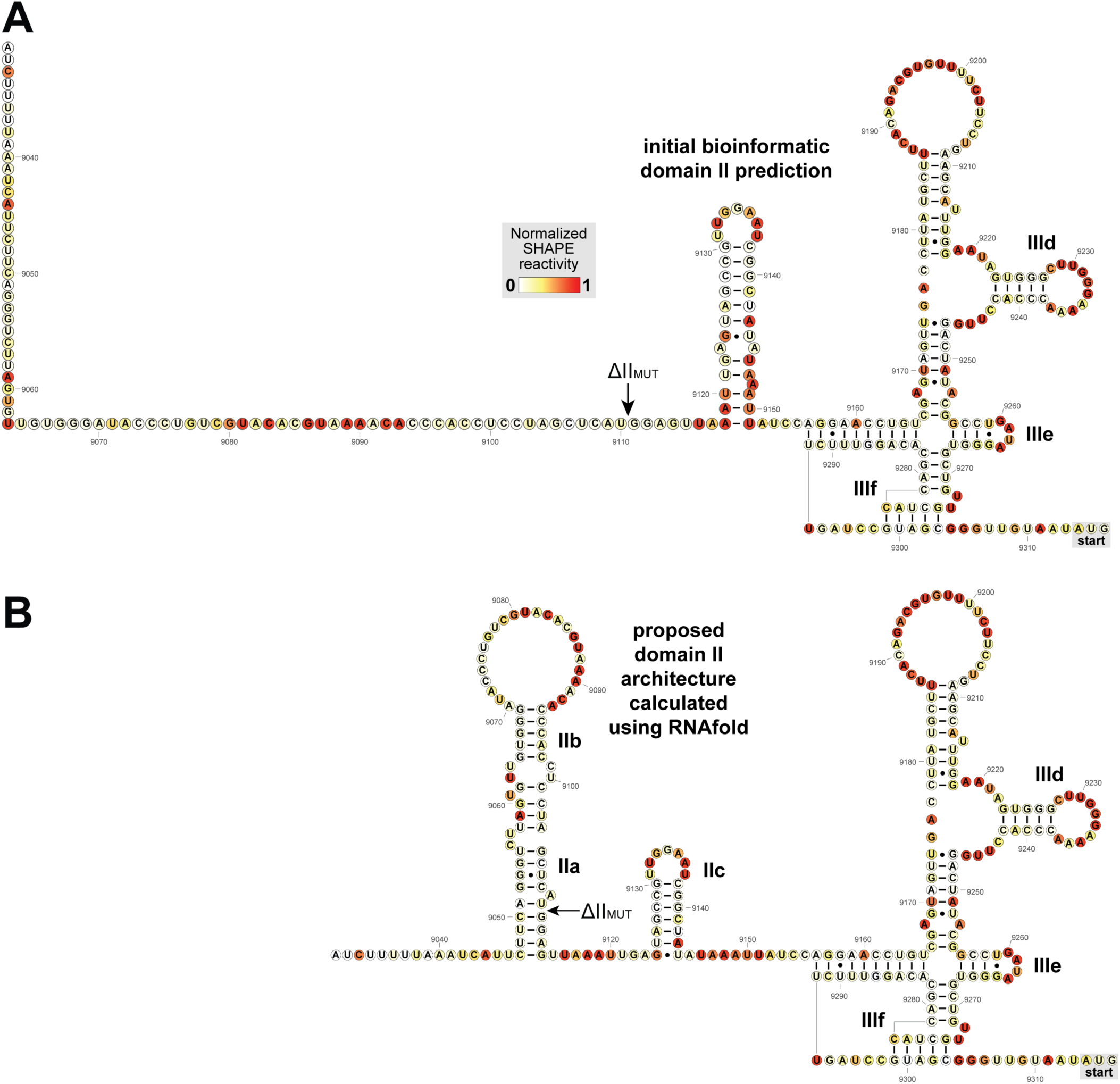
Chemical probing data for the MeV-E 3’ IRES mapped onto different domain II secondary structure models. (A,B) Normalized reactivity at each nucleotide position of the MeV-E 3’ IRES RNA, starting directly after the ORF1 stop codon, to the chemical probing reagent 1-methyl-7-nitroisatoic anhydride (1M7). Reactivities are mapped onto the small stem loop structure predicted by the bioinformatic alignment output from Infernal (panel A) or our proposed secondary structure model of its domain II equivalent (panel B)

**Figure S3.**
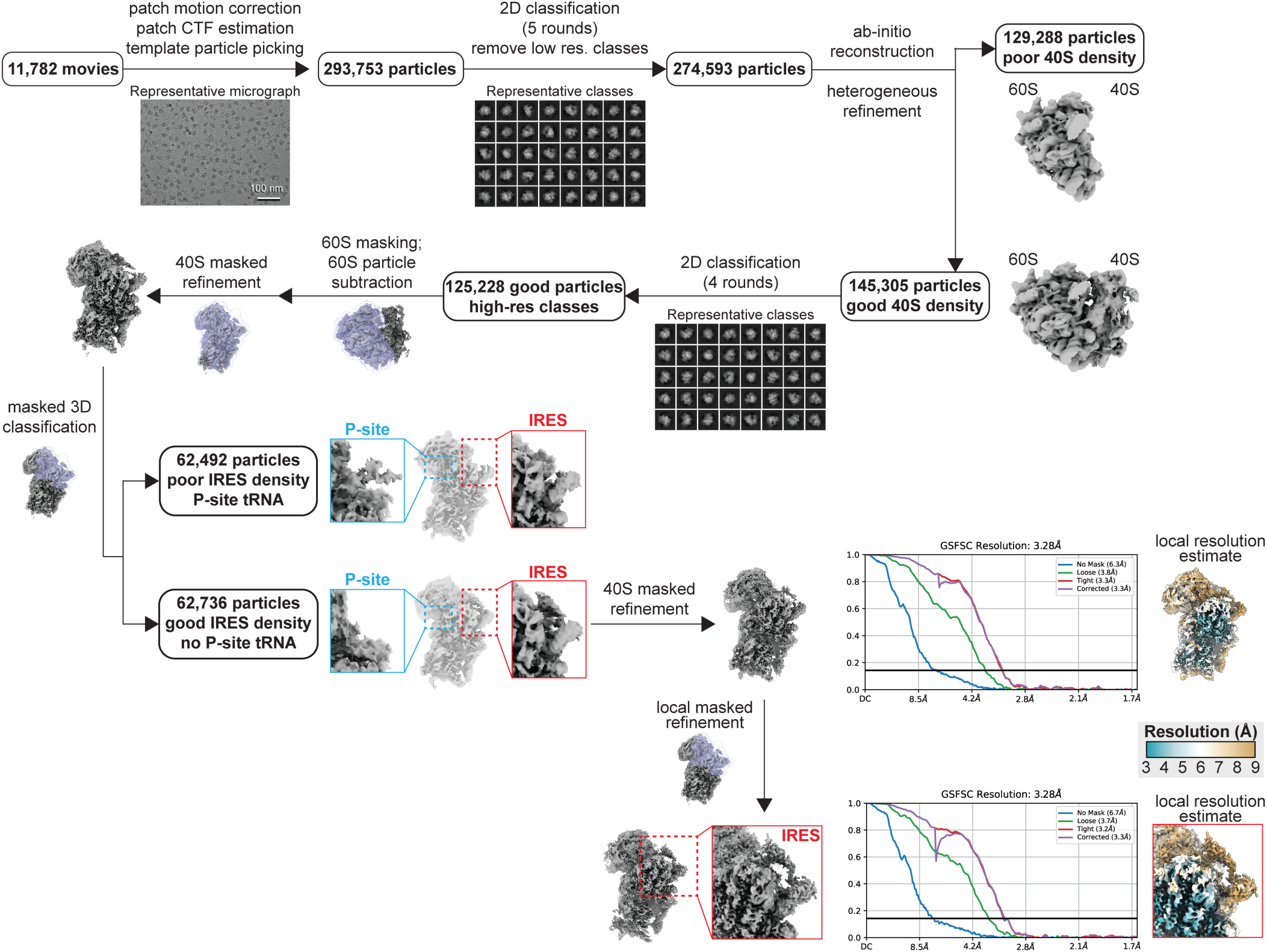
Overview of the data processing workflow for single particle cryoEM analysis of the MeV-E 3’ IRES RNA-80S structure. The cryoEM processing pipeline in cryoSPARC for the MeV-E 3’ IRES RNA-80S complex using data collected with a 300 kV electron microscope. After particle picking and several rounds of 2D classification to remove junk particles and low-resolution classes, ab initio reconstruction followed by heterogeneous refinement were performed to remove particles with poor 40S density. After local motion correction, the 60S was masked (purple) and particle subtracted. 3D classification of remaining 40S particles led to identification and removal of an undesired state containing tRNA and poorer IRES density. The final 40S map (3.28 Å resolution) was generated using 62,736 particles in a non-uniform refinement, and the final map of the masked IRES region (purple) was generated from the same set of particles with a local refinement. Estimation of overall resolution of cryoEM maps using the gold standard FSC of 0.143.

**Figure S4.**
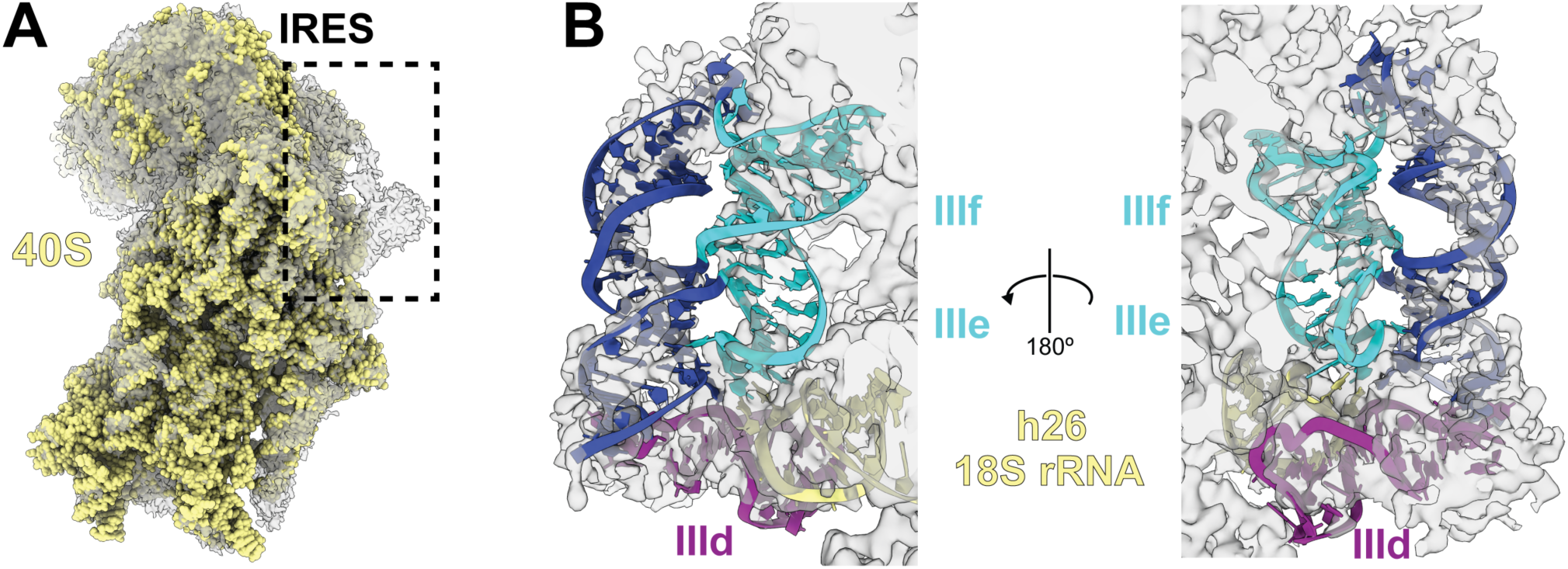
Refined and focused cryoEM maps of the MeV-E 3’ IRES RNA-40S structure with models fit into the density. (A) CryoEM map of the MeV-E 3’ IRES-ribosome complex fit with an apo-40S structure to demonstrate the area of the map (boxed) assigned to the IRES RNA. (B) Two views of the model of the 40S-bound MeV-E 3’ IRES RNA structure with its binding site on the apical portion of helix 26 (h26, also called expansion segment 7) of the 18S rRNA (yellow) fit into the cryoEM map of the MeV-E 3’ IRES-ribosome complex.

**Figure S5.**
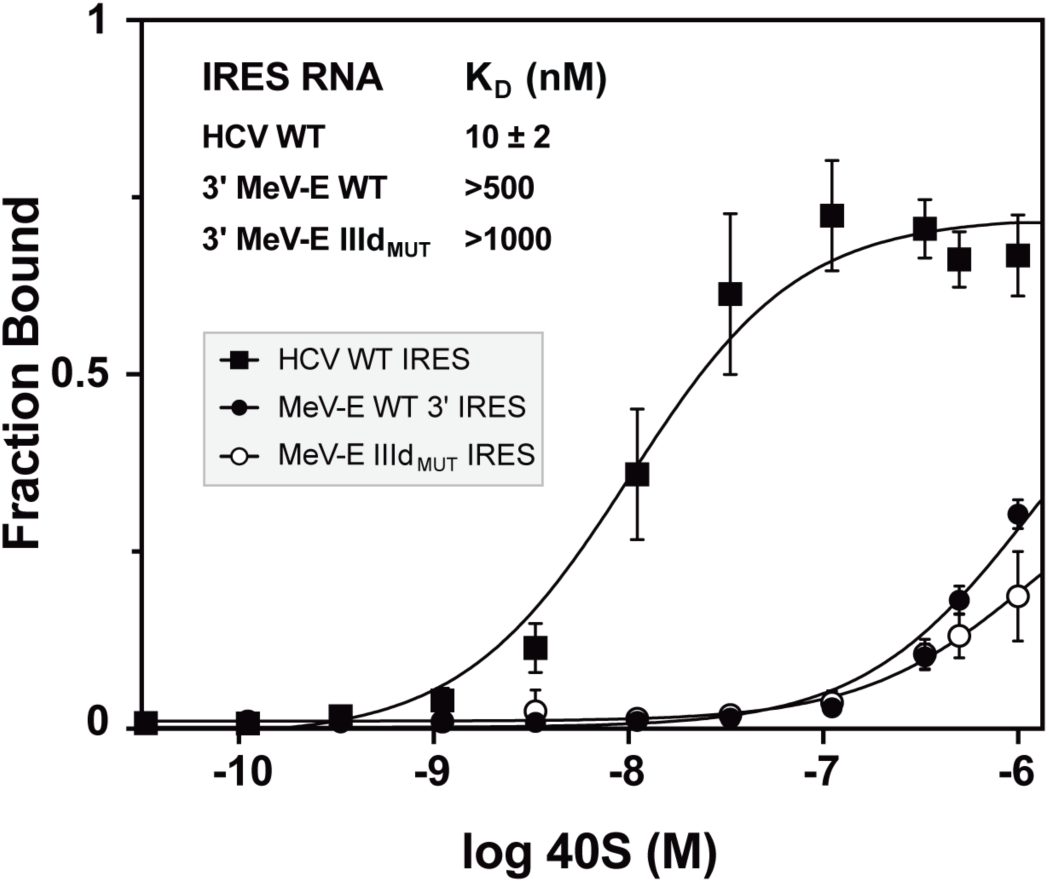
40S binding affinities of the HCV and MeV-E 3’ IRESs. Dissociation coefficients (K_D_) of IRES RNAs for purified 40S subunits as determined by *in vitro* filter binding experiments. Data plotted are the average and standard deviation of three separate replicates and fit with a 1:1 sigmoidal dose response curve. Values for the MeV-E 3’ IRES constructs are estimated as binding is not saturated at 1 μM.

**Figure S6.**
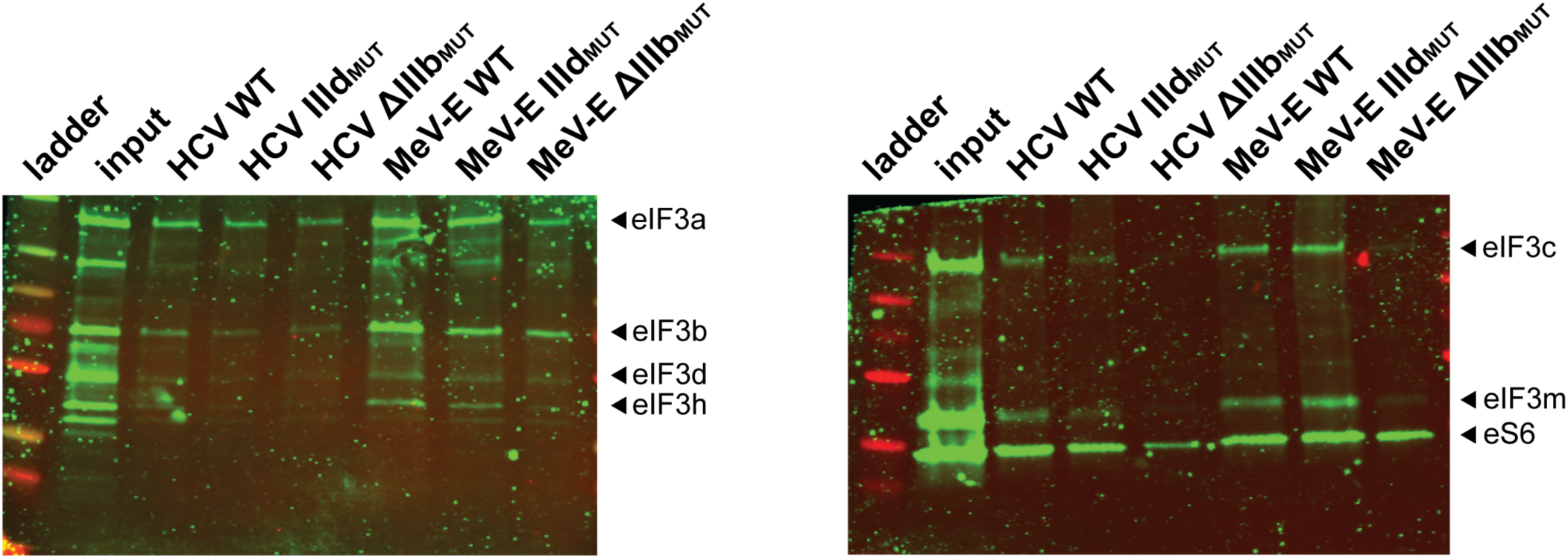
Presence of eIF3 in IRES preinitiation complexes. Western blots of the input and samples pulled out of lysate with biotinylated IRES RNAs using primary antibodies against components of the eIF3 complex (eIF3a,b,c,d,h,m) and a 40S ribosomal protein control (eS6). Data shown are identical samples split between two blots and are representative of three independent experiments.

**Table S1.**
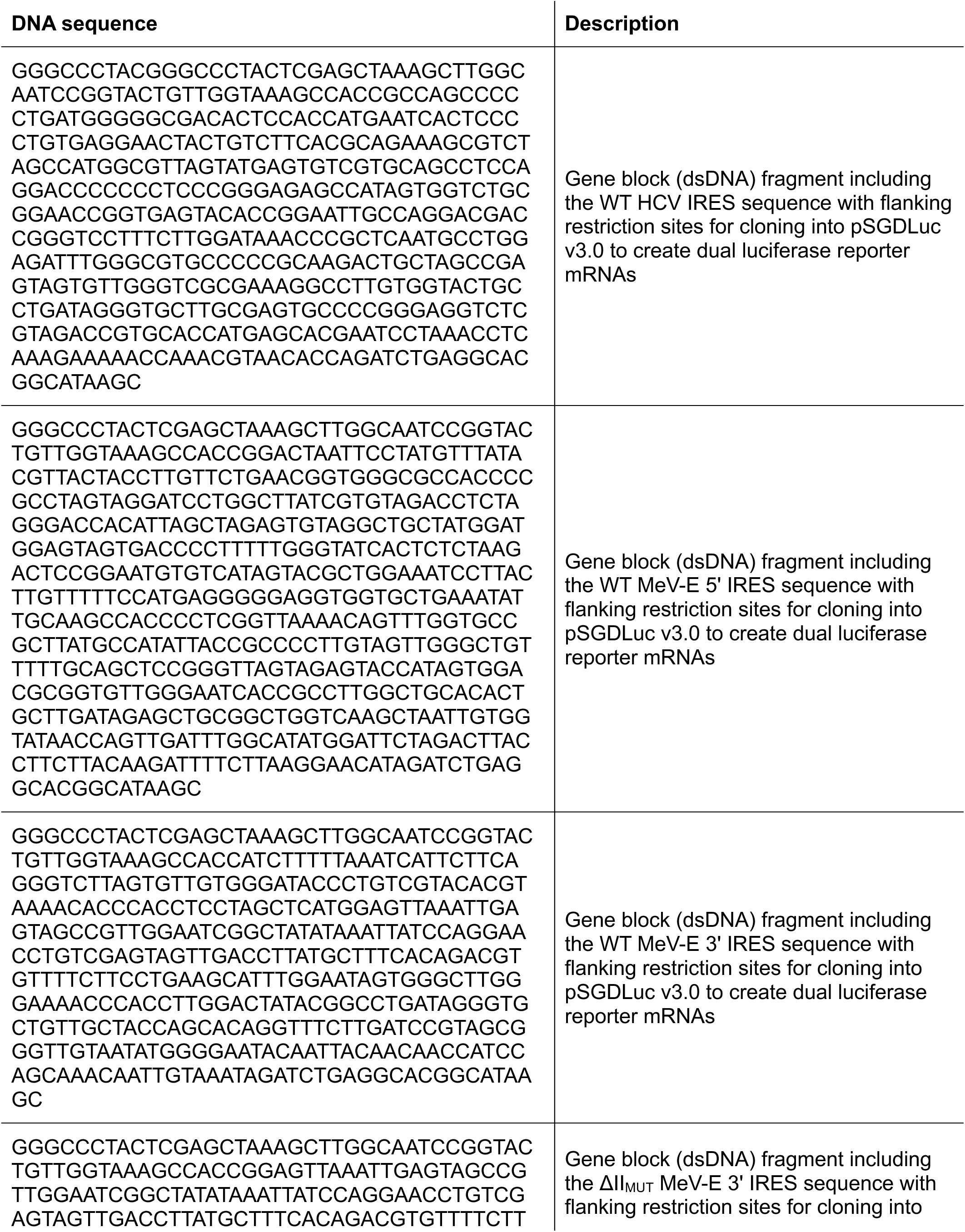
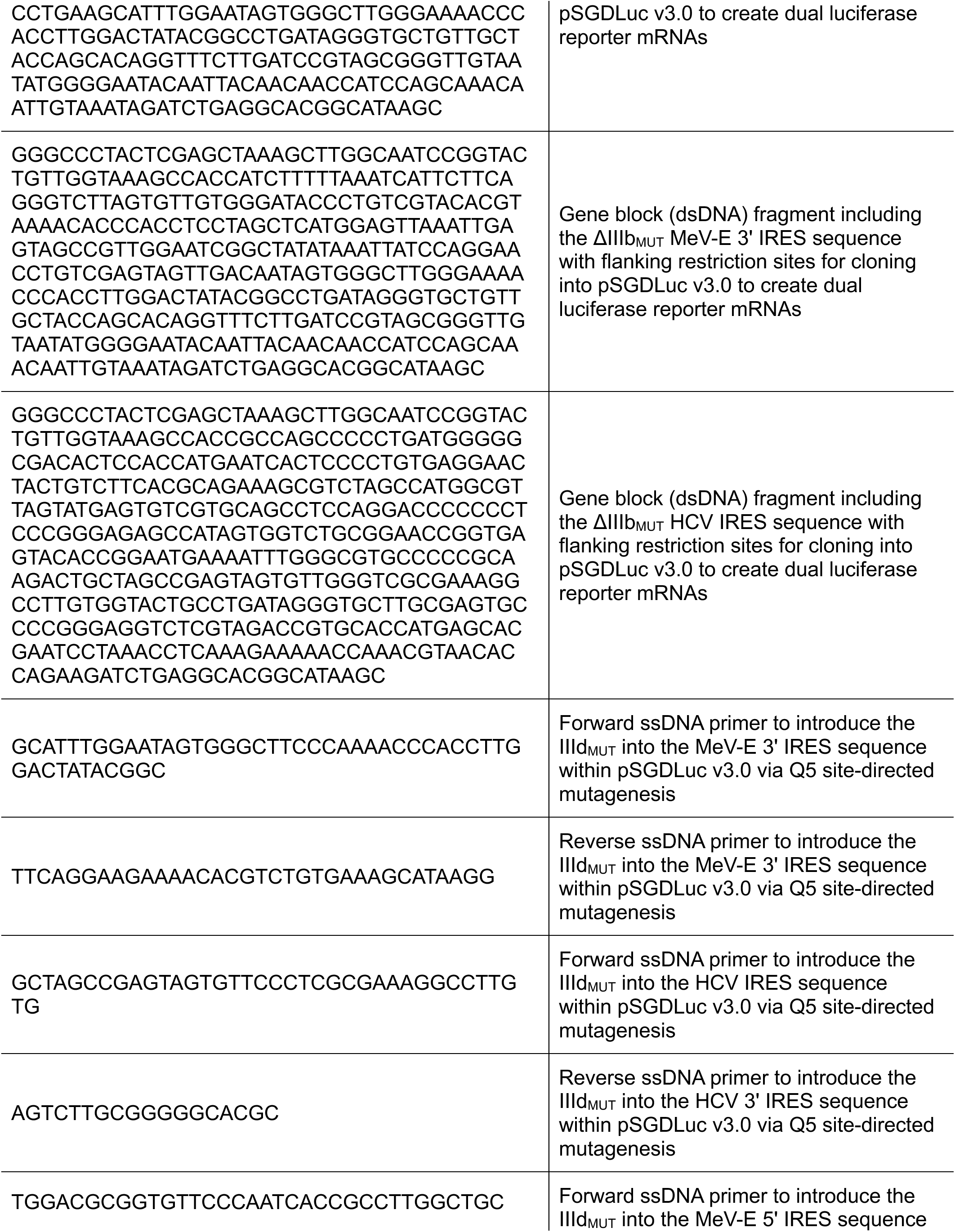
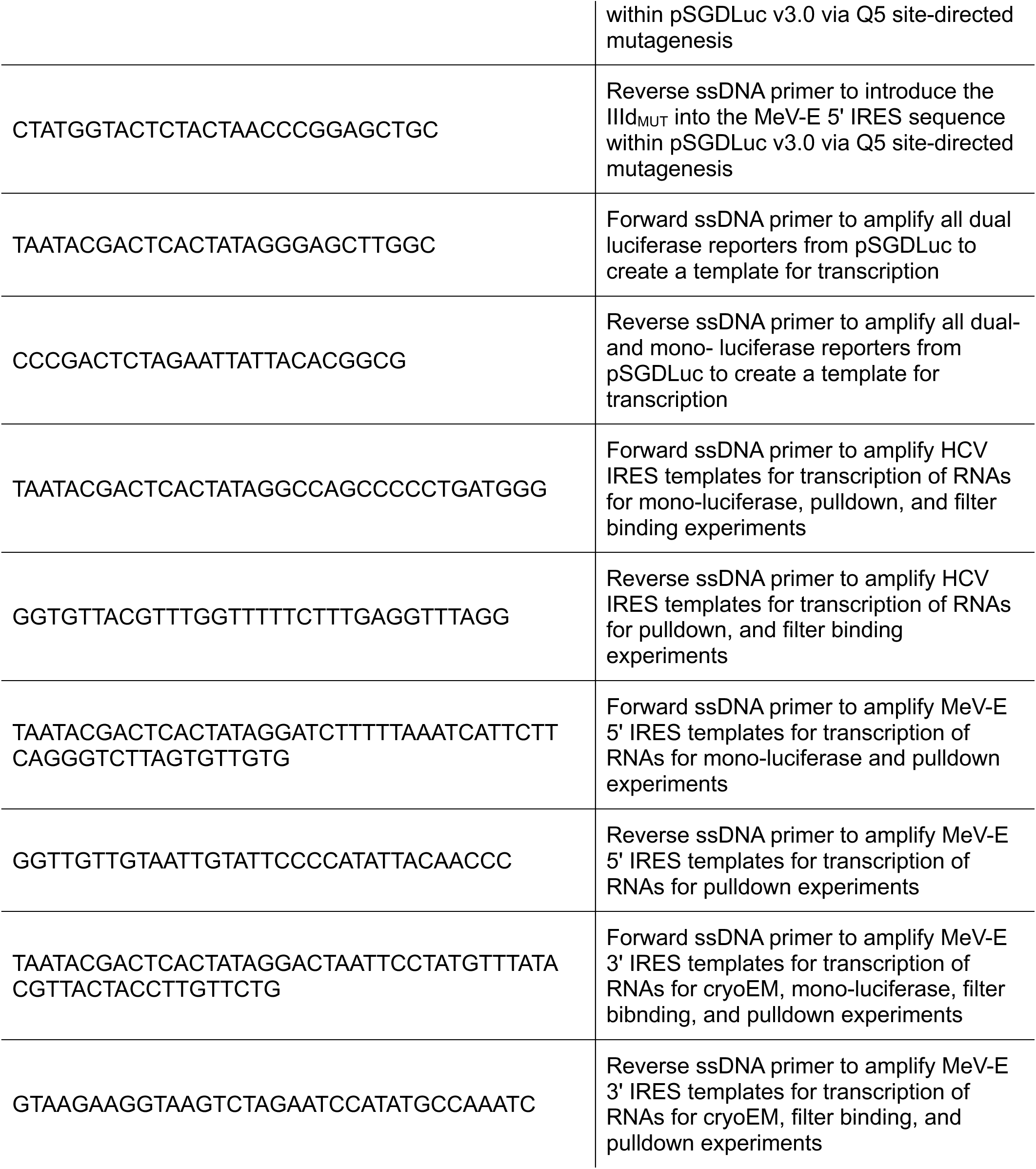
DNA oligonucleotide sequences used in this study.

**Table S2.** Cryo-EM data collection, refinement, and validation statistics.

| **MeV-E IRES RNA-80S** |  |
| --- | --- |
| **Data collection and processing** |  |
| Microscope | Titan Krios |
| Voltage (KeV) | 300 |
| Camera | Gatan K3 |
| Magnification | 105,000 |
| Pixel size (Å) | 0.8255 |
| Total electron exposure (e-/Å<sup>2</sup>) | 50 |
| Defocus range (µm) | -0.4 to -1.8 |
| Tilt angle | 0°, 30° |
| Energy filter | BioContinuum |
| Energy filter slit width (eV) | 20 |
| Micrographs collected | 11,782 |
| Total extracted particles (no.) | 293,753 |
| **MeV IRES RNA-40S** |  |
| **Reconstruction** |  |
| Final refined particles | 64,736 |
| Resolution, FSC threshold 0.143 (Å) | 3.28 |
| Resolution range, FSC threshold 0.05 (Å)<br>min/median/max | 1.752/8.596/57.121 |
| Map sharpening B factor (Å<sup>2</sup>) | -25 |
| **Model refinement** |  |
| Initial models used (PDB code) | 5A2Q, 7SYP |
| Model composition |  |
| Chains | 36 |
| Non-H Atoms | 77,254 |
| Residues Protein | 4896 |
| Residues Nucleotides | 1787 |
| Bond (RMSD) |  |
| Lengths (Å) | 0.004 |
| Angles (°) | 0.652 |
| MolProbity score | 2.20 |
| Clash Score | 18.08 |
| Ramachandran Plot (%) |  |
| Outliers | 0.02 |
| Allowed | 6.92 |
| Favored | 93.06 |
| Rama-Z (Z-score, RMSD) |  |
| Whole | -1.53 |
| Helix | -0.05 |
| Sheet | -1.02 |
| Loop | -1.59 |
| Rotamer Outliers (%) | 0.00 |
| B factors (Å<sup>2</sup>) min/max/mean |  |
| Nucleotide | 6.46/236.57/100.12 |
| Protein | 5.25/162.48/73.13 |

**Table S3.** Genomic information for megrivirus sequences.

| **Virus name** |  | **Abbreviation** | **NCBI genome accession number** |
| --- | --- | --- | --- |
| turkey hepatitis virus | melegrivirus A | MeV-A1 | KF961188.1 |
| duck megrivirus | megrivirus A2 | MeV-A2 | KC663628 |
| goose megrivirus | megrivirus A3 | MeV-A3 | KY369299 |
| pigeon mesivirus 1 | megrivirus B1 | MeV-B1 | KC876003 |
| pigeon mesivirus 2 | megrivirus B2 | MeV-B2 | KC811837 |
| chicken megrivirus | megrivirus C1 | MeV-C1 | KF961186 |
|  | megrivirus C2 | MeV-C2 | KF979336 |
|  | megrivirus C3 | MeV-C3 | MG846465 |
| harrier megrivirus | megrivirus D1 | MeV-D | KY488458 |
| penguin megrivirus | megrivirus E1 | MeV-E | MF405436 |

